# The epithelial-mesenchymal transcription factor *SNAI1* represses transcription of the tumor suppressor miRNA *let-7* in cancer

**DOI:** 10.1101/2020.11.04.368688

**Authors:** H Wang, E Chirshev, N Hojo, T Suzuki, A Bertucci, M Pierce, C Perry, R Wang, J Zink, CA Glackin, YJ Ioffe, JJ Unternaehrer

## Abstract

We aimed to determine the mechanism of epithelial-mesenchymal transition (EMT)-induced stemness in cancer cells. Cancer relapse and metastasis are caused by rare stem-like cells within tumors. Studies of stem cell reprogramming have linked *let-7* repression and acquisition of stemness with the EMT factor, *SNAI1*. The mechanisms for the loss of *let-7* in cancer cells are incompletely understood. In four carcinoma cell lines from breast cancer, pancreatic cancer and ovarian cancer and in ovarian cancer patient-derived cells, we analyzed stem cell phenotype and tumor growth via mRNA, miRNA, and protein expression, spheroid formation, and growth in patient-derived xenografts. We show that treatment with EMT-promoting growth factors or *SNAI1* overexpression increased stemness and reduced *let-7* expression, while *SNAI1* knockdown reduced stemness and restored *let-7* expression. Rescue experiments demonstrate that the pro-stemness effects of *SNAI1* are mediated via *let-7. In vivo*, nanoparticle-delivered siRNA successfully knocked down *SNAI1* in orthotopic patient-derived xenografts, accompanied by reduced stemness and increased *let-7* expression, and reduced tumor burden. Chromatin immunoprecipitation demonstrated that *SNAI1* binds the promoters of various *let-7* family members, and luciferase assays revealed that *SNAI1* represses *let-7* transcription. In conclusion, the *SNAI1/let-7* axis is an important component of stemness pathways in cancer cells, and this study provides a rationale for future work examining this axis as a potential target for cancer stem cell-specific therapies.

**Novelty and Impact:** This study provides new insight into molecular mechanisms by which EMT transcription factor *SNAI1* exerts its pro-stemness effects in cancer cells, demonstrating its potential as a stem cell-directed target for therapy. *In vitro* and *in vivo*, mesoporous silica nanoparticle-mediated *SNAI1* knockdown resulted in restoration of let-7 miRNA, inhibiting stemness and reducing tumor burden. Our studies validate *in vivo* nanoparticle-delivered RNAi targeting the *SNAI1/let-7* axis as a clinically relevant approach.

## Introduction

Cancer stem-like cells (CSC) are the subpopulation of tumor cells responsible for long-term maintenance of tumors. These cells are capable of self-renewal and differentiation, making them an important contributor to tumor recurrence^1^. The origin of CSC is not completely understood. In some cancers, normal tissue stem cells appear to be altered to result in CSC^1–4^, while in others, somatic cells appear to be reprogrammed to the stem cell fate^5–7^. Whether the cells of origin in carcinomas are tissue resident stem cells or reprogrammed somatic cells, some aspects of the process by which CSC attain stem cell features are comparable to somatic cell reprogramming^4,7–9^. In somatic cell reprogramming, cells lose their differentiated characteristics and take on an embryonic or stem cell phenotype. Similarly, stem cells in tumors dedifferentiate and express genes consistent with the oncofetal state^10–12^.

An important factor in maintenance of the differentiated state is the tumor suppressor miRNA *let-7. Let-7*, consisting in humans of nine highly conserved members in eight chromosomal locations, plays crucial roles in differentiation^13^. Because the seed sequence of the individual family members is identical, and the remaining sequence is different at only 1-3 residues, this miRNA family is generally presented as having redundant roles^13^. In pluripotent cells and germ cells, miRNA *let-7* expression is low, while differentiated cells uniformly express high levels^14^. Factors required for stemness (a property referring to a cell’s ability to self-renew and differentiate^3^) are inhibited by *let-7*^15^. Loss of *let-7* is thus necessary for the stem cell state, either in reprogramming or in cancer^13,16,17^. *Let-7* represses a set of embryonic genes and oncogenes, and its loss allows upregulation of those genes, resulting in the oncofetal state^13–16^. Replacing *let-7* reduces the stem cell population and reduces resistance to chemotherapy^18^. These data strongly implicate *let-7* as a key regulator of the CSC phenotype.

*Let-7* is frequently reduced in many types of cancer^13^. *Let-7* loss correlates with poor prognosis, and is a biomarker for less differentiated cancer^13,19^, and predicts tumor growth and metastasis^20^. Mechanisms for its loss are incompletely understood. miRNAs are regulated transcriptionally, epigenetically, and post-transcriptionally^21^. The pluripotency-associated factor *LIN28A* blocks *let-7* biogenesis by inhibiting its processing to the mature form, but *LIN28A* is absent in differentiated cells^22^. Several factors have been shown to regulate *let-7* transcription, including the epithelial-mesenchymal transition (EMT) transcription factor TWIST1, TP53, MYC, BMI1, NFKB1, and CEBPA^18^. We set out to study *let-7* regulation at the transcriptional level, because of evidence for its importance in dedifferentiation^17^ and potential influence on the metastatic disease course.

EMT is a fundamental process for development and homeostasis whereby epithelial cells lose their cell polarity and cell-cell adhesion, and gain the migratory and invasive features typical of mesenchymal cells^23^. The aberrant activation of EMT is considered to be a hallmark of cancer metastasis^23–25^. Many studies have found that EMT is not an all-or-none response; instead, it is a multi-step process, with cells existing in states ranging from fully epithelial to fully mesenchymal. Cells are observed in several intermediate or partial (hybrid) EMT states^25^. In fact, cancer cells that undergo partial EMT (cells without complete loss of epithelial morphology or complete acquisition of mesenchymal morphology) have been reported to pose a higher metastatic risk^26,27^. Besides metastasis, cancer cells that undergo EMT demonstrate enhanced stemness, including tumor initiation ability and capacity to differentiate to multiple lineages^2,28,29^. The subpopulation within cancer cells that have higher stemness has been shown to contribute to the tumor’s invasiveness and resistance to therapies^30,31^. Hence, targeting stem-like cancer cells via EMT may be a crucial step to improve patient outcome.

Much evidence connects EMT with the acquisition of stem cell properties. Cells that have undergone EMT acquire the ability to differentiate to multiple lineages^28^. The expression of EMT transcription factors *SNAI1, TWIST1*, or *ZEB1* results in an increase in the proportion of cells with stem cell properties^29,32,33^. Recent work demonstrates that it is the cells with hybrid properties that are most active in assays for stemness, and *SNAI1* expression locks cells in the hybrid state^34^. *SNAI1’s* transcription factor roles include repression of epithelial factors such as *CDH1*, stimulation of mesenchymal factors, and repression of miRNAs such as miR-34^23,35^. We chose to focus on the EMT factor *SNAI1* because of its role in reprogramming somatic cells to pluripotency^17,36^ and in cancer stemness^29,33,37^. Furthermore, *SNAI1* interacts with the miRNA of interest – *let-7*. It binds *let-7* family promoters and its early upregulation in reprogramming correlates with loss of *let-7*^17^. Because the increase of *SNAI1* and the decrease of *let-7* occurred at time points in reprogramming prior to upregulation of *LIN28A*, we hypothesized that it might be the loss of *let-7*, rather than the gain of *LIN28A*, that destabilized the differentiated state. In the studies presented here, we asked whether these reprogramming principles applied in cancer: does expression of *SNAI1* lead to loss of *let-7* and gain of stemness?

One promising approach to target genes such as EMT transcription factors is antisense oligonucleotide strategies, but avoiding degradation by ubiquitous nucleases, preventing immune activation, and allowing extravasation and cellular uptake to targeted cells present technical challenges for this technique^38^. Poor cytoplasmic delivery of RNA therapeutics to appropriate cells has inhibited research progress, but our team has optimized a targeted nanoparticle delivery method to deliver RNAis to tumors^39^. Mesoporous silica nanoparticles (MSN) are small (50-200nm), but have relatively large surface area due to their pore structure^40^. Coating them with cationic polyethylenimine (PEI) facilitates loading of siRNA cargo, and conjugation with hyaluronic acid (HA) assists in delivery to target cells^41,42^: HA is the ligand for CD44, enriched on the surface of ovarian cancer stem cells^43^.

In this study, we hypothesized that *SNAI1* directly represses miRNA *let-7* transcription, and that *SNAI1* knockdown would result in restoration of *let-7* expression and reduction of stemness and tumor growth. Using breast, pancreatic, and ovarian cancer cells, transforming growth factor beta-1 (TGFB1) or epidermal growth factor (EGF) treatment or *SNAI1* overexpression increased stemness and reduced *let-7* expression, while *SNAI1* knockdown reduced stemness and increased *let-7* expression. We demonstrate on the molecular level that *SNAI1* binds promoters of *let-7* family members in cancer cells. Luciferase assays demonstrate that the presence of *SNAI1* reduces *let-7* transcription, consistent with direct repression of *let-7* by *SNAI1*. Thus, one mechanism by which EMT promotes stemness is via loss of *let-7*, destabilizing the differentiated state. With the utilization of the orthotopic patient-derived xenograft (PDX) murine models of high grade serous ovarian carcinoma (HGSOC), we demonstrate feasibility of *in vivo SNAI1* knockdown, delivering siRNA with mesoporous silica nanoparticles. In orthotopic PDX, *SNAI1* knockdown results in increased *let-7* levels and reduced tumor growth.

## Materials and Methods

### Cell cultures

The human HGSOC cell line OVSAHO (RRID:CVCL_3114) was the kind gift of Gottfried Konecny (University of California Los Angeles), and OVCAR8 (RRID:CVCL_1629) was from Carlotta Glackin (City of Hope). HEK293T (RRID:CVCL_0063), PANC-1 (RRID:CVCL_0480) (gift of Nathan Wall, Loma Linda University (LLU)), MCF-7 (RRID:CVCL_0031) (gift of Eileen Brantley, LLU), OVSAHO and OVCAR8 cells were cultured in Dulbecco’s Modification of Eagle’s Medium (DMEM) with 10% fetal bovine serum (FBS), 2mM of L-Glutamine, 100 U/mL of penicillin, and 10 μg/mL of streptomycin. NCCIT (RRID:CVCL_1451), used as a positive control for expression of pluripotency factors, was cultured in RPMI with 10% FBS, 2mM L-Glutamine, 1mM sodium pyruvate, 100 U/mL of penicillin, and 10 μg/mL of streptomycin. MCF-7 and PANC-1 cells were treated with TGFB1 (10 ng/ml), OVCAR8 and OVSAHO cells were treated with EGF (100 ng/ml). PDX6, a HGSOC chemotherapy naïve sample, was obtained as described^20^. All studies were approved by the Loma Linda University (LLU) institutional review board (IRB). Deidentified fresh ovarian cancer ascites samples was provided by the LLU Biospecimen Laboratory and were processed by centrifuging. Erythrocytes were removed by overlaying a cell suspension on a 3ml Ficoll gradient. Cells were initially engrafted into NSG mice subcutaneously in the region of the mammary fat pad, resulting in PDX. Patient-derived samples were cultured in three parts Ham’s F12 and one part DMEM, supplemented with 5% FBS, 10uM insulin, 0.4uM hydrocortisone, 2 ug/ml isoprenaline, 24 ug/ml adenine, 100 U/ml of penicillin, and 10 ug/mL streptomycin. 5-10 uM Y27632 was added to establish growth *in vitro*^44^. Low passage (maximal passage number: 15) patient-derived cells were used to avoid changes induced by extensive passaging in *in vitro* culture.

All human cell lines have been authenticated using STR profiling within the last three years. All experiments were performed with mycoplasma-free cells.

### Reverse-transcription quantitative PCR (RT-qPCR)

Total RNA from cell culture samples was isolated using TRIzol reagent (Life Technologies, Carlsbad, CA, USA) according to the manufacturer’s instructions. For mRNA expression analysis, cDNA was synthesized with 1 μg of total RNA using Maxima First Strand cDNA Synthesis Kit (K1672; Thermo fisher scientific, Grand Island, NY, USA). Real-time RT-qPCR for mRNA was performed using PowerUP SYBR Green master mix (Thermo fisher scientific, Grand Island, NY, USA) and specific primers on a Stratagene Mx3005P instrument (Agilent Technology, Santa Clara, CA, USA). Primer sequences are listed in Supplementary Table 2. For miRNA expression analysis, cDNA was synthesized with 100 ng of total RNA using specific stem-loop RT primers and TaqMan microRNA Reverse Transcription Kit (Applied Biosystems, Foster City, CA, USA). Real-time RT-qPCR for miRNA was performed using TaqMan Universal PCR Master Mix II (Applied Biosystems, Foster City, CA, USA) with specific probes (Life Technologies 4440887 assay numbers 000377 (let-a), 002406 (let-7e), 002282 (let-7g), 002221 (let-7i), U47 (001223)) on a Stratagene Mx3005P instrument (Agilent Technology, Santa Clara, CA, USA). The results were analysed using the ΔΔ cycles to threshold (ΔΔCt) method; ACTB (mRNA) and U47 (miRNA) were used for normalization.

### Western blot

Proteins were extracted from cells in PBS by adding SDS sample buffer (2% SDS, 2.5% beta-mercaptoethanol, 7.5% glycerol) and then sonicated for 10 - 15 sec. 30 ul of lysate per sample (2.4×10e5 cells) were heated to 100°C for 5 min and then loaded on SDS-PAGE gel [4-12%]. After running at 150 V for 20-40 min, samples were transferred to PVDF membrane. Membranes were incubated in 5% milk for blocking for 1 hr at room temperature. After blocking and washing with 1X TBST, membranes were incubated in primary antibodies diluted at the appropriate dilution (as suggested by manufacturer data sheets) over night at 4°C. Antibodies used include: HMGA2 (D1A7, Cell Signaling Technology, Danvers, MA), GAPDH (14C10, Cell Signaling Technology, Danvers, MA), SNAI1 (L70G2; Cell Signaling Technology, Danvers, MA), α/β-TUBULIN (2148S; Cell Signaling Technology, Danvers, MA, USA). Secondary antibody incubations were done with an anti-mouse IgG conjugated with DyLight 800 (SA5-10176; Invitrogen, Carlsbad, CA, USA) or anti-rabbit IgG antibody conjugated with DyLight 680 (35569; Invitrogen, Carlsbad, CA, USA) at 1/30000 for 1 hr at room temperature. Immunoblots were scanned and visualized using Odyssey Infrared Imaging System (LI-COR Biosciences, Lincoln, NE, USA). Densitometry was performed on scanned immunoblots by ImageJ software (National Institutes of Health, Bethesda, MD, USA). Quantification of Western blot data was done by measuring the intensity of bands of the protein of interest divided by the intensity of the samples’ own α/β-TUBULIN bands (ImageJ).

### Retroviral overexpression

The cDNA of human *SNAI1* was subcloned from Flag-Snail WT (Addgene 16218) into pWZL-Blast-GFP (Addgene 12269) after removing GFP using BamH1/Xho1. Retroviral particles were produced in HEK293T cells after co-transfection of retrovirus plasmid vector pWZL-Blast-Flag-Snail or control vector pWZL-Blast-Flag-Empty with packaging plasmids (VSVG, Gag/pol) using polyethylenimine (PEI) (Polyscience). After 48h and 72h, supernatant containing virus was collected and filtered through a 0.22 μM filter. Supernatants were used for cell transduction or stored at −80 °C. Cells were transduced with retrovirus in the presence of 6 μg/ml protamine sulfate and selected with 5 ug/ml Blasticidin (InvivoGen #ant-bl-05) for 5 days.

### DsiRNA-mediated knockdown

A panel of dicer-substrate small inhibitory RNAs (DsiRNA, IDT) were screened for SNAI1 knockdown (Supplementary Figure 3). HA-conjugated, PEI-coated MSNs were synthesized as described^41^; in brief, MSNs were produced using the sol-gel method, dissolving 250mg cetyltrimethylammonium bromide in 120ml water with 875ul of 2M sodium hydroxide solution. Second, 1.2ml tetraethylorthosilicate was added, stirred for 2 hours, allowing formation of MSN. Particles were collected by centrifugation, and washed with methanol and acidic methanol. Low molecular weight cationic PEI (1.8 kDa branched polymer) was electrostatically attached to the MSN surface to provide a positive charge to attract negatively charged siRNA^41^, and HA was covalently bound to the amine groups in the PEI using EDC-NHS coupling reaction^45^. DsiRNA targeting SNAI1 or control (oligonucleotides sequence listed in Supplementary Table 4) were used for knockdown *in vitro*, loaded on MSN as described^39^. To complex siRNA for *in vitro* experiments, 10 μl siRNA at 10 μM was mixed with 70 μl MSNs at 500 μg/ml and 20 μl water, and the mixture was incubated overnight at 4 °C on a rotor. The following day, 100 μl of the HA-MSN-siRNA complexes was added to each well of a 6-well plate containing 1900 μl normal medium. To complex siRNA for *in vivo* experiments, 15 μl siRNA at 10 μM was mixed with 105 μl HA-MSNs at 500 μg/ml, and the mixture was incubated overnight at 4 °C on a rotor. The following day, 120 μl of the HA-MSN-siRNA complexes were injected intravenously (tail vein). For *in vivo* experiments, HA-MSN-siRNA were injected twice weekly.

### Mimic transfection

Let-7i mimics (sense: 5’ - mCmArGmCrAmCrAmArAmCrUmArCmUrAmCrCmUrCA - 3’; antisense 5’ - /5Phos/rUrGrArGrGrUrArGrUrArGrUrUrUrGrUrGrCrUmGmUrU - 3’) and scrambled control mimics (sense 5’ - mCmArUmArUmUrGmCrGmCrGmUrAmUrAmGrUmCrGC - 3’; antisense5’ - /5Phos/rGrCrGrArCrUrArUrArCrGrCrGrCrArArUrArUmGmG rU - 3’; IDT) were reverse transfected at 2nM using Lipofectamine RNAiMax (Life Technologies) according to manufacturer guidelines.

### Chromatin Immunoprecipitation (ChIP)

ChIP assay was conducted using MAGnify™ Chromatin Immunoprecipitation System (Thermo Fisher Scientific, #49-2024) according to manufacturer directions. Untreated OVCAR8, OVSAHO, MCF-7 cells with or without 10 ng/mL of TGFB1 were crosslinked with 1% formaldehyde. 1.25 M glycine in cold PBS were then added to stop the crosslinking reaction. Cell lysates were prepared with lysis buffer with protease inhibitors (50μL per 1 million cells). Chromatin was then sheared into 200-500-bp fragments using *Fisher Scientific Sonic Dismembrator Model F60 With Probe*. Each immunoprecipitation (IP) reaction contains 100,000 cells. Dynabeads^®^ were coupled with anti-Snail (L70G2; Cell Signaling Technology, Danvers, MA) or Mouse IgG (supplied in MAGnify kit) as negative controls (1 μg per CHIP). After 1 hour on a rotor, these antibody-Dynabeads^®^ complexes were incubated with chromatin and put on rotor for 2 hours at 4°C. As input control, 10 μL of diluted chromatin were put aside without binding to the antibody-Dynabeads^®^ complexes. After chromatin-Antibody-Dynabeads^®^ complexes were washed with IP Buffer to remove unbound chromatin. Reverse Crosslinking Buffer was added to reverse the formaldehyde crosslinking. Real-time RT-qPCR for DNA was performed using PowerUP SYBR Green master mix (Thermo fisher scientific, Grand Island, NY, USA) and specific primers on a Stratagene Mx3005P instrument (Agilent Technology, Santa Clara, CA, USA). Primer sequences are listed in Supplementary Table 3. The results were analyzed using the ΔΔ cycles to threshold (ΔΔCt) method; ACTB was used for normalization.

### Luciferase assays

HEK293T cells were plated at 50,000 cells per well. Twenty-four hours later PEI reagent was used to transfect cells with 200ng full length *let-7*, truncated *let-7i* (*lucB*), or mutated *let-7i* (*mlucB*) promoter luciferase vector in combination with 5ng Renilla luciferase, and 200ng *SNAI1*-expressing or empty vector (Addgene 16218). Forty-eight hours post transfection (or twenty-four hours for promoter truncation/mutation) dual-luciferase reporter assay kit (Promega) was used to analyze bioluminescence on SpectraMax i3x microplate reader (Molecular Devices, Sunnyvale, CA, USA). *Let-7a1df1* promoter luciferase was a kind gift from Dr. Zifeng Wang^46^, *let-7a3* from Dr. Hillary Coller^47^, *Let-7c* from Dr. Maria Rizzo^48,49^, full length *let-7i* from Dr. Steve O’Hara^50^, and truncated (lucB)/mutated (mlucB) let-7i from Dr. Muh-Hwa Yang^51^.

### Spheroid formation assay

Cells were plated at a density of 10,000 cells/mL (12,000 cells/ml for PDX6 cells) in non-tissue culture coated plates, 10 technical replicates per condition, and maintained in serum-free medium (DMEM/F12 50/50) supplemented with 0.4% bovine serum albumin, 10ng/mL FGF, 20ng/mL EGF, 6.7ng/ml selenium, 5.5ug/ml transferrin, 10ug/ml insulin, and 1% knock out serum replacement (Gibco/ThermoFisher Scientific) for 7 days. Secondary spheroid assays were done by harvesting after seven days, trypsinization, and re-seeding at 10,000 cells/mL, followed by seven additional days of growth. To determine the number and size of spheroids, phase contrast images of spheroids taken on a Nikon Eclipse Ti microscope were analyzed using ImageJ software (National Institutes of Health, Bethesda, MD, USA).

### Mice

All animal procedures were conducted according to animal care guidelines approved by the Institutional Animal Care and Use Committee at Loma Linda University. Orthotopic PDX experiments were carried out in nude mice (nu/nu), obtained from Jackson Laboratory (Sacramento, CA, USA), which were housed in specific pathogen-free conditions, and were used for xenografts at 6-10 weeks of age.

### Orthotopic xenograft model and live animal imaging

To allow *in vivo* visualization, PDX6 cells were transduced with a CMV-p:EGFP-ffluc pHIV7 lentiviral vector (eGFP-ffluc, kind gift of Christine Brown)^52^, which encodes a fusion protein of GFP and firefly luciferase. The eGFP-ffluc-transduced PDX6 cells were selectively isolated based on GFP expression via FACSAria cell sorter (BD Biosciences, San Jose, CA, USA). PDX6 cells were injected into the right ovarian bursa of nude mice with Matrigel (354248; Corning, Corning, NY, USA) at 2.5×10^5^ cells per mouse, eight mice per condition. For *in vivo* experiments, DsiRNA with 2’-O-methyl modifications were used^53^ (oligonucleotides sequence listed in Supplementary Table 4). Starting 1 week after initial injection and continuing twice weekly, HA-MSN-siRNA were injected intravenously. After intraperitoneal injection of luciferin, the mice were imaged with an IVIS Lumina Series III *in vivo* imaging system (PerkinElmer, Waltham, MA, USA). Live imaging was performed twice weekly and the bioluminescent images were analyzed using Living Image *in vivo* Imaging Software (PerkinElmer, Waltham, MA, USA) to assess tumor burden at primary and metastatic sites. At day 1, 16 mice were randomized and assigned into two groups (siControl and siSnail, 8 mice each). The bioluminescence of animals from each group was measured at each time point. Based on tumor development, some mice were censored from analyses. Each animal’s measurement was normalized to its own bioluminescence from day one and then the means for each time point were analyzed using a two-way ANOVA. To determine endpoints, mouse abdominal girth was measured prior to surgery and monitored once a week. When the first mouse reached the endpoint of an increase of 25% in girth, all mice were euthanized, and necropsy was carried out. Primary and metastatic tumor weight and tumor locations were recorded, and samples were harvested for gene and protein expression analysis.

### Statistical analyses

For all *in vitro* experiments, cell samples in the same treatment group were harvested from at least 3 biological replicates and processed individually. For *in vivo* experiments, data are from one representative experiment of three. All values in the figures and text are the means ± SD. Statistical analyses were performed using the Prism 7.0a for Mac OS X (GraphPad Software, Inc.). Statistical significance among mean values was determined by Student’s t-test with two-tailed alpha level of 0.05 considered significant, with the exception of tumor growth in the *in vivo* study, which is determined by two-way ANOVA with Tukey’s multiple comparison test. *, P < 0.05; **, P < 0.01; ***, P < 0.001; ****, P < 0.0001.

## Results

### *SNAI1* leads to increased stemness

To test the relationship between *SNAI1* expression and changes in stemness, we induced *SNAI1* expression with growth factors including TGFB1 and EGF^54,55^. We tested several cancer cell lines of epithelial origin including pancreatic (PANC-1), breast (MCF-7), and ovarian (OVCAR8 and OVSAHO).

After two days of TGFB1 (MCF-7 or PANC-1) or EGF (OVSAHO or OVCAR8) treatment, as expected, RNA and protein expression levels of *SNAI1* increased, confirmed by RT-qPCR (Figure 1A) and Western blot (Figure 1C, Supplementary Figure 2A). TGFB1 does not induce *SNAI1* expression in OVSAHO or OVCAR8 (Supplementary Figure 1B); for this reason, ovarian cancer cell lines were treated with EGF. The smaller change in SNAI1 protein observed in OVCAR8 could be explained by its high endogenous *SNAI1* level as previously described^56^; endogenous levels of all cell lines are shown in Supplementary Figure 1A. mRNA expression of stemness markers *LIN28A, NANOG, POU5F1* and *HMGA2* increased after treatment (Figure 1B). Western blot analysis showed an increase of HMGA2 protein in OVSAHO (43%) (Figure 1D, Supplementary Figure 2B); however, this was not detectable in other lines. We used spheroid assays as a measure of self-renewal and growth in non-adherent conditions, which are increased with higher stemness^56,57^. In agreement with the phenotypic measurements above, cells in which *SNAI1* expression was induced via TGFB1 or EGF formed more spheroids, indicating a higher frequency of cells with stem cell attributes (Figure 1E). This trend is more significant after one passage, where the increase in number of spheroids formed upon TGFB1 or EGF treatment is even larger (Supplementary Figure 5B). Along with the increased *SNAI1* expression, consistent with a change to a more stem cell-like gene expression pattern, we observed a decrease in expression levels of *let-7* family members (Figure 1F). We chose to follow one *let-7* member from each of four clusters on chromosomes 3, 9, 12, and 19^21^.

**Figure 1.**
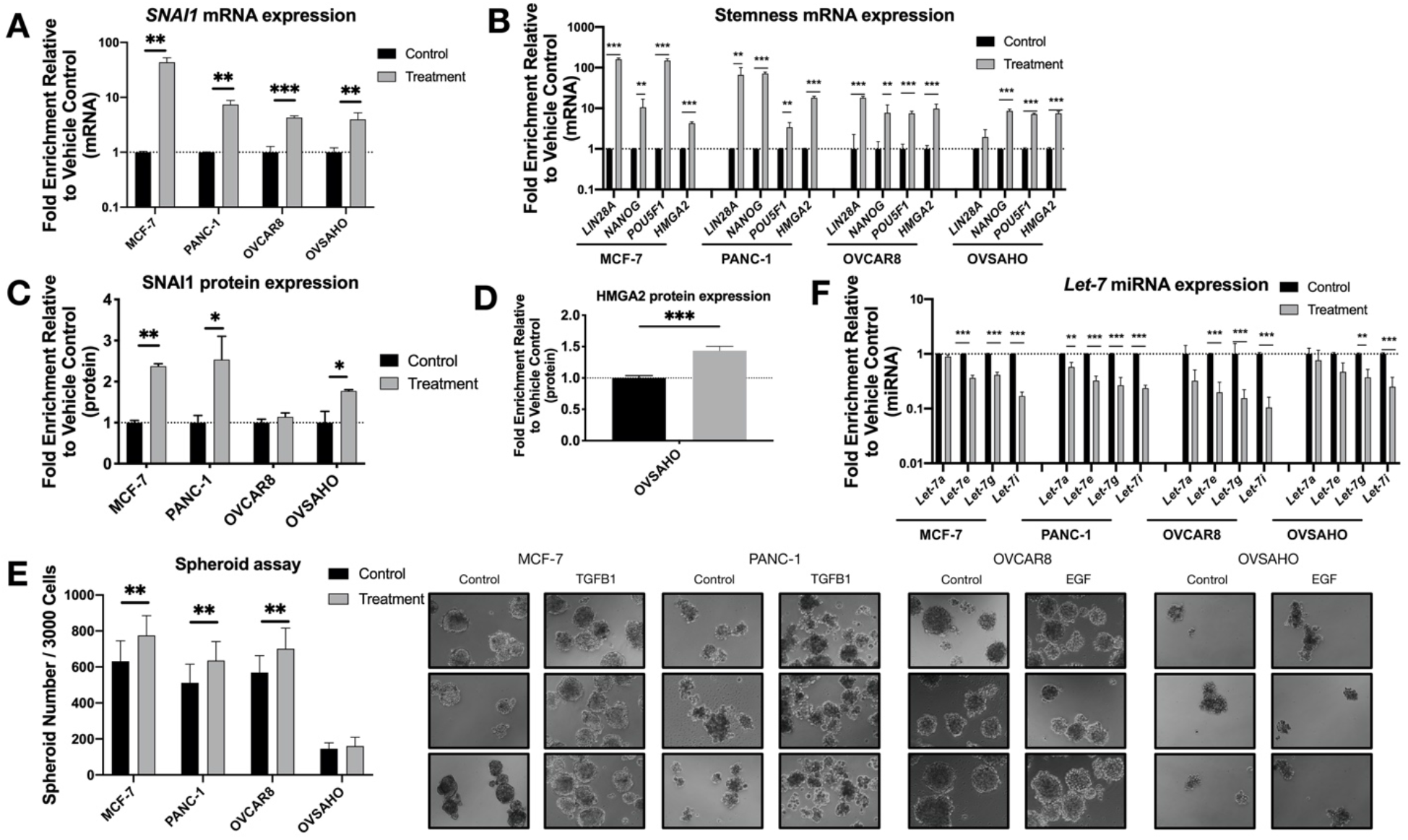
Growth factor treatment results in increased SNAI1, stemness and decreased *let-7* expression. MCF-7, PANC-1 were treated with TGFB1; OVCAR8, OVSAHO were treated with EGF. Levels of control group (cells treated with vehicle control) were normalized to 1; note that values for RT-qPCR are shown on a log scale. A,B. RT-qPCR analysis for mRNA expression level of *SNAI1* (A) and stemness markers (B, *LIN28A, NANOG, POU5F1* and *HMGA2*). C,D. The quantification of Western blot analysis for protein expression of SNAI1 (C) and HMGA2 (D). E. Left panel: the quantification of number of spheroids per 3000 cells is shown. Right panel: Phase contrast images of spheroids formed from cells as indicated are presented; in each panel, the spheroids formed from control group are presented on the left, those from the treatment group are on the right. F. RT-qPCR analysis for *let-7* miRNA (*let-7a, let-7e, let-7g* and *let-7i*) expression.

Because growth factor-induced EMT resulted in changes consistent with an increase in stemness, we wished to pinpoint mechanisms of stemness downstream of EMT. Our previous studies indicated a role for *SNAI1* in the induction of the stem cell fate^17^. Besides inducing EMT, the TGFB1 signaling pathway is important in mediating cellular proliferation, preventing progression through the cell cycle, and multiple other actions^54^. EGF also plays an important role in the development of tumors by regulating cell proliferation, differentiation, migration and angiogenesis^55^. Thus, treatment with these growth factors changes the expression of numerous genes besides *SNAI1*. To specify the effect of a single factor, *SNAI1*, we overexpressed *SNAI1* to determine whether it alone could induce the stem cell state. Cell lines were virally transduced with constitutively expressed *SNAI1* or control vector.

After transduction, the increase in *SNAI1* mRNA and protein expression (Figure 2A, 2C and Supplementary Figure 4A) was accompanied by a significant increase in stemness markers *LIN28A, POU5F1*, and *HMGA2* (Figure 2B). Western blot data confirmed this change, showing an increase in HMGA2 (Figure 2D, Supplementary Figure 4B). With the increase in expression of *SNAI1* and stemness genes, we observed a decrease in *let-7* family members (Figure 2F). Consistent with the phenotypic changes, *SNAI1* overexpression led to an increase in the number of spheroids formed (Figure 2E, Supplementary Figure 4C) (the size of spheroids for OVCAR8 is quantified and presented in Supplementary Figure 5A), to a greater extent in secondary spheroids (Supplementary Figure 5B), suggesting increased stemness associated with *SNAI1*. In order to investigate whether the regulation of stemness is directly through SNAI1’s action on *let-7*, we overexpressed *let-7i* in *SNAI1* overexpressing cells (Supplementary Figure 6A). *Let-7i* overexpression resulted in abrogation of *SNAI1*-induced stemness as measured by RT-qPCR (Supplementary Figure 6B) and spheroid formation (Supplementary Figure 6C, D). These results confirmed that *SNAI1* overexpression is sufficient to shift the phenotype toward stemness via its effect on *let-7*.

**Figure 2.**
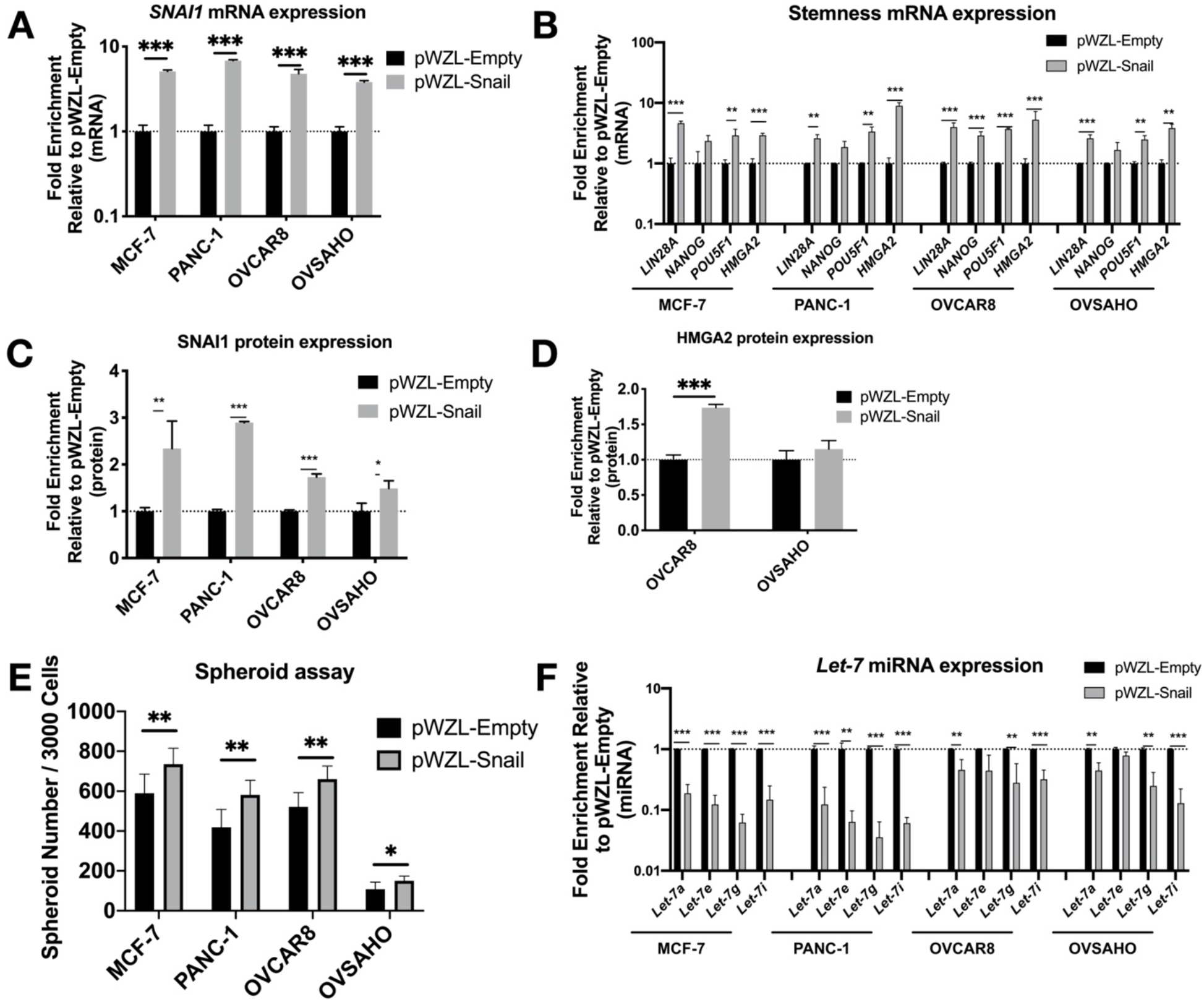
*SNAI1* overexpression results in increased stemness and decreased *let-7* expression. Cell lines were transduced with the retroviral expression vector pWZL-Snail or empty vector, pWZL-Empty in cell lines MCF-7, PANC-1, OVCAR8 and OVSAHO. Levels of control group (cells transduced with pWZL-Empty) were normalized to 1; note that values for RT-qPCR are shown on a log scale. A,B. RT-qPCR analysis for mRNA expression of *SNAI1* (A) and stemness markers (B, *LIN28A, NANOG, POU5F1*, and *HMGA2*). C,D. The quantification of Western blot analysis for protein expression of SNAI1 (C) and HMGA2 (D). E. The quantification of number of spheroids formed per 3000 cells as indicated. F. RT-qPCR analysis for *let-7* miRNA (*let-7a, let-7e, let-7g* and *let-7i*) expression.

### *SNAI1* knockdown reverses stemness

Having established the impact of *SNAI1’s* gain-of-function on cells’ stemness and *let-7* levels, we proceeded to knock down *SNAI1* to test if the opposite effects could be observed. We used HA-conjugated MSN^41^ (HA-MSN) to deliver siRNA in MCF-7, PANC-1, OVSAHO and OVCAR8. We observed a decrease in the mRNA expression level of *SNAI1* after HA-MSN-siSnail treatment in most cases (Figure 3A). The knockdown of *SNAI1* was confirmed on the protein level with Western blot data (Figure 3C, Supplementary Figure 7A). Together with the decrease of *SNAI1*, the expression of stemness markers also decreased on the mRNA level (Figure 3B). HMGA2 protein also decreased in PANC-1 and OVSAHO after siSnail treatment (Figure 3D, Supplementary Figure 7B). *SNAI1* knockdown resulted in reduced frequency of stem cells, as measured by number of spheroids formed (Figure 3E, Supplementary Figure 7C), and secondary spheroids showed a greater difference between siSnail and siControl (Supplementary Figure 5B). An increase in spheroid size in OVCAR8 was also observed (Supplementary Figure 5A). Consistent with the *SNAI1* time course, *let-7* expression increased after *SNAI1* knockdown (Figure 3F). Similar effects were observed with two siRNAs (Supplementary Figure 8). These results indicate that reducing *SNAI1* expression leads to decreased stemness as well as restoration of *let-7* expression in cancer cells.

**Figure 3.**
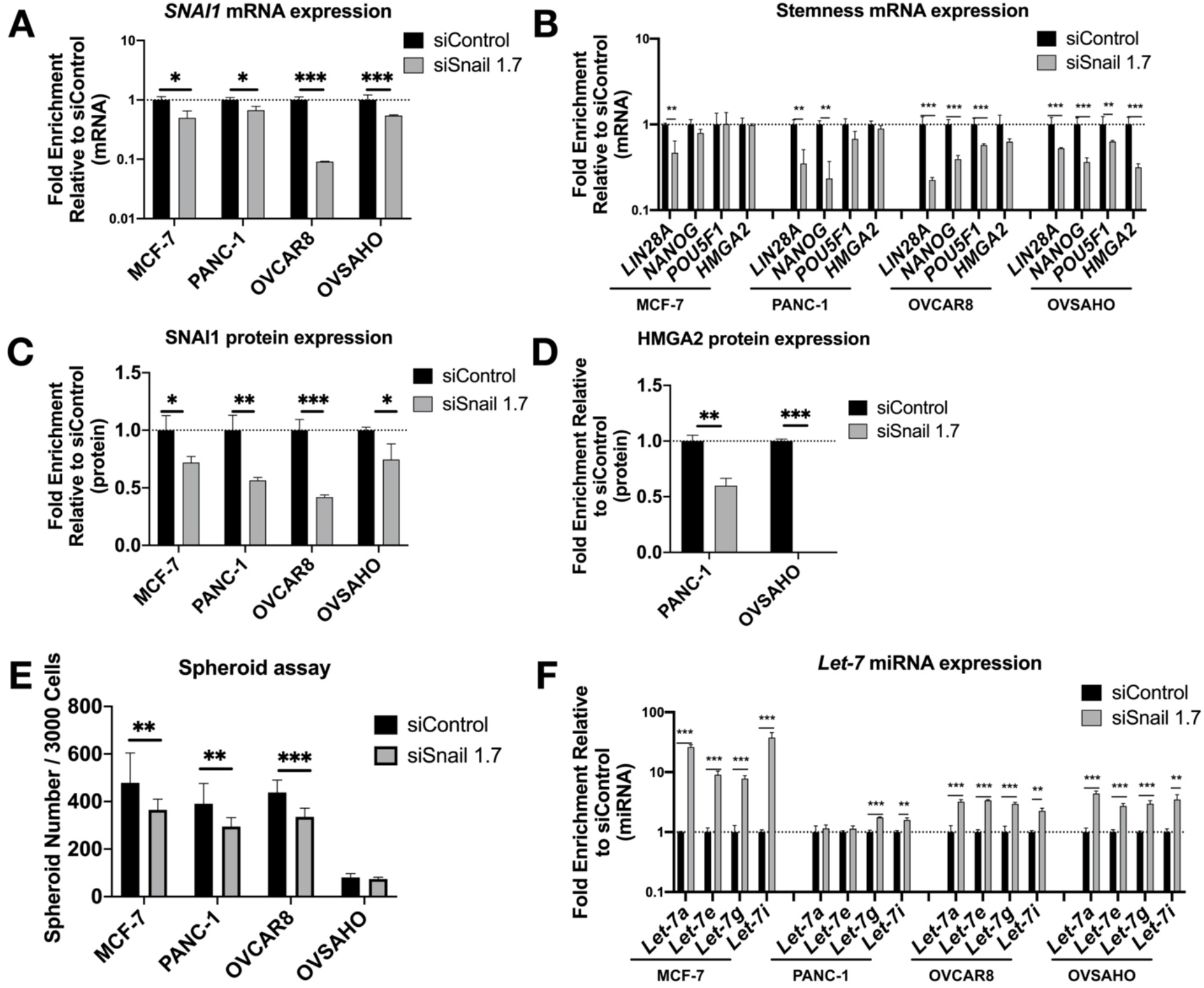
*SNAI1* knockdown reverses stemness and restores *let-7* expression. Mesoporous silica nanoparticles coated with hyaluronic acid (HA-MSN) were used to deliver siRNA (siSnail and siControl) in MCF-7, PANC-1, OVCAR8 and OVSAHO. Levels of control group (cells treated with siControl) were normalized to 1; note that values for RT-qPCR are shown on a log scale. A,B. RT-qPCR analysis for mRNA expression of *SNAI1* (A) and stemness markers (B, *LIN28A, NANOG, POU5F1*, and *HMGA2*). C,D. The quantification of Western blot analysis for protein expression of SNAI1 (C) and HMGA2 (D). E. The quantification of number of spheroids formed per 3000 cells as indicated. F. RT-qPCR analysis for *let-7* miRNA (*let-7a, let-7e, let-7g* and *let-7i*) expression.

### *SNAI1* knockdown reverses stemness in patient derived HGSOC samples *in vitro* and decreases tumor burden *in vivo*

To test our findings in a more clinically relevant setting, we knocked down *SNAI1* in patient-derived cells *in vitro* using HA-MSN-siSnail (Figure 4A, C and Supplementary Figure 9A). In agreement with our observations in cell lines, PDX cells treated with HA-MSN-siSnail showed decreased levels of stemness markers (Figure 4B,D and Supplementary Figure 9A), decreased size (Supplementary Figure 5A) and number of spheroids formed (Figure 4E, Supplementary Figure 9B), and increased levels of *let-7* (Figure 4F).

**Figure 4.**
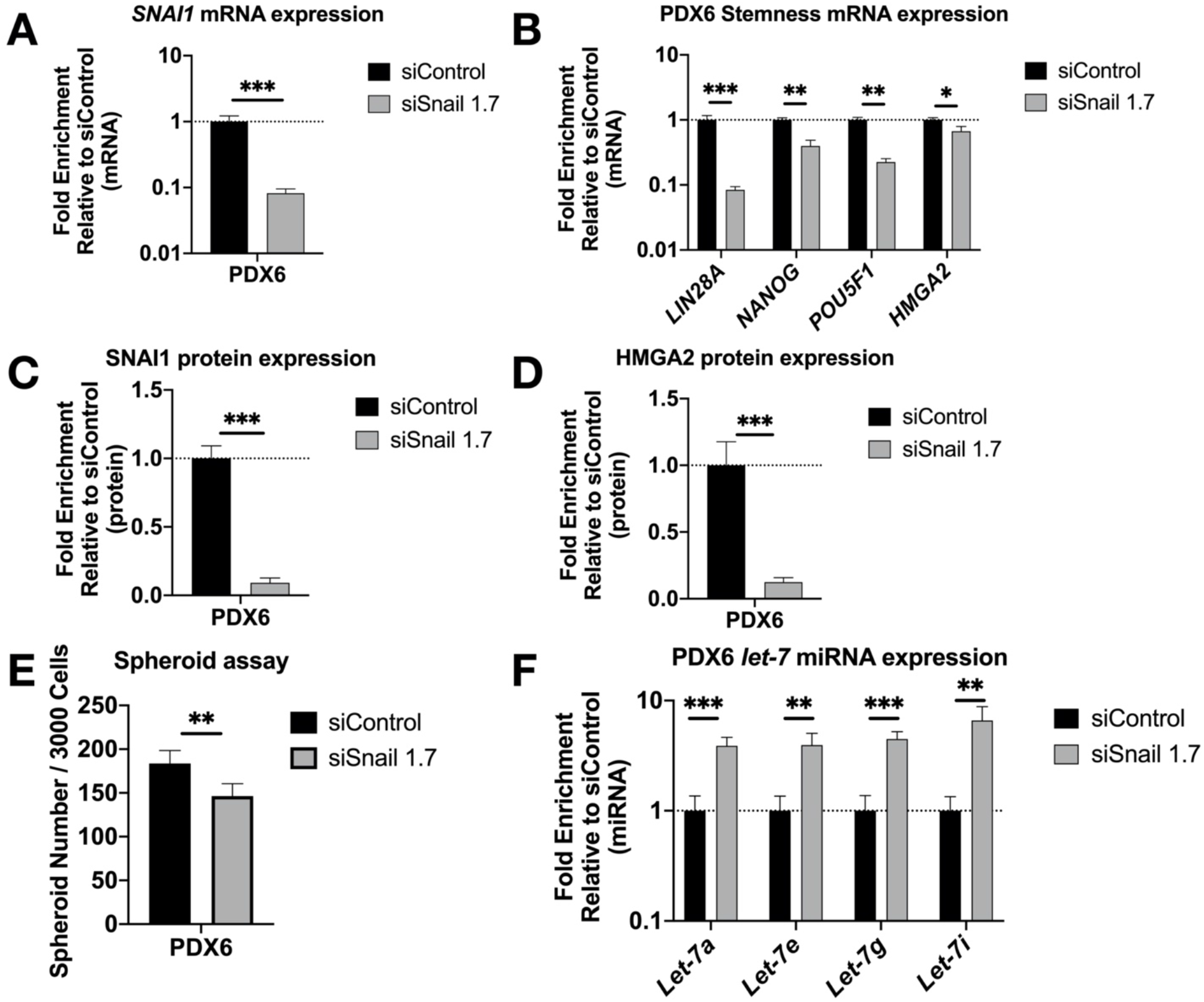
*SNAI1* knockdown reduces stemness in patient-derived cells *in vitro*. HA-MSN were used to deliver siRNA (siSnail and siControl) in PDX cells *in vitro*. Levels of control group (cells treated with siControl) were normalized to 1; note that values for RT-qPCR and Western blot are shown on a log scale. A,B. RT-qPCR analysis for mRNA expression of *SNAI1* (A) and stemness markers (B, *LIN28A, NANOG, POU5F1*, and *HMGA2*). C,D. The quantification of Western blot analysis for protein expression of SNAI1 (C) and HMGA2 (D). E. The quantification of number of spheroids per 3000 cells formed from PDX6 *in vitro*. F. RT-qPCR analysis for *let-7* miRNA (*let-7a, let-7e, let-7g* and *let-7i*) expression.

To extend these results to an *in vivo* setting, luciferized PDX6 cells were injected into the ovarian bursa of nude mice in our orthotopic xenograft model^56^. Mice were imaged twice weekly for bioluminescence, and total flux was quantified over seven weeks. One week after bursa injection, treatment with HA-MSN-siSnail (or HA-MSN-siControl) began and continued twice weekly for the duration of the experiment. Upon necropsy, RT-qPCR results showed a decrease of *SNAI1* along with reduced *LIN28A, NANOG and POU5F1* in tumors from mice treated with HA-MSN-siSnail (Figure 5A, B). In agreement with mRNA results, the protein levels of SNAI1, LIN28A and HMGA2 were significantly decreased in mice treated with HA-MSN-siSnail (Figure 5C, D and Supplementary Figure 10A). Consistent with the *in vitro* results, *let-7* levels were also increased in mice treated with HA-MSN-siSnail (Figure 5E). In addition, primary tumor weights demonstrated smaller tumors in siSnail mice (Supplementary Figure 10B). Visualization of tumors in live animals revealed that primary tumors were significantly smaller in mice receiving HA-MSN-siSnail injections (Figure 5F). These results demonstrate that *SNAI1* was successfully knocked down *in vivo* using targeted nanoparticle-delivered RNAi. Taken together, our results demonstrate that knockdown of *SNAI1* in patient derived HGSOC samples *in vitro* and *in vivo* results in restoration of *let-7*, decreased stemness, and reduced tumor burden.

**Figure 5.**
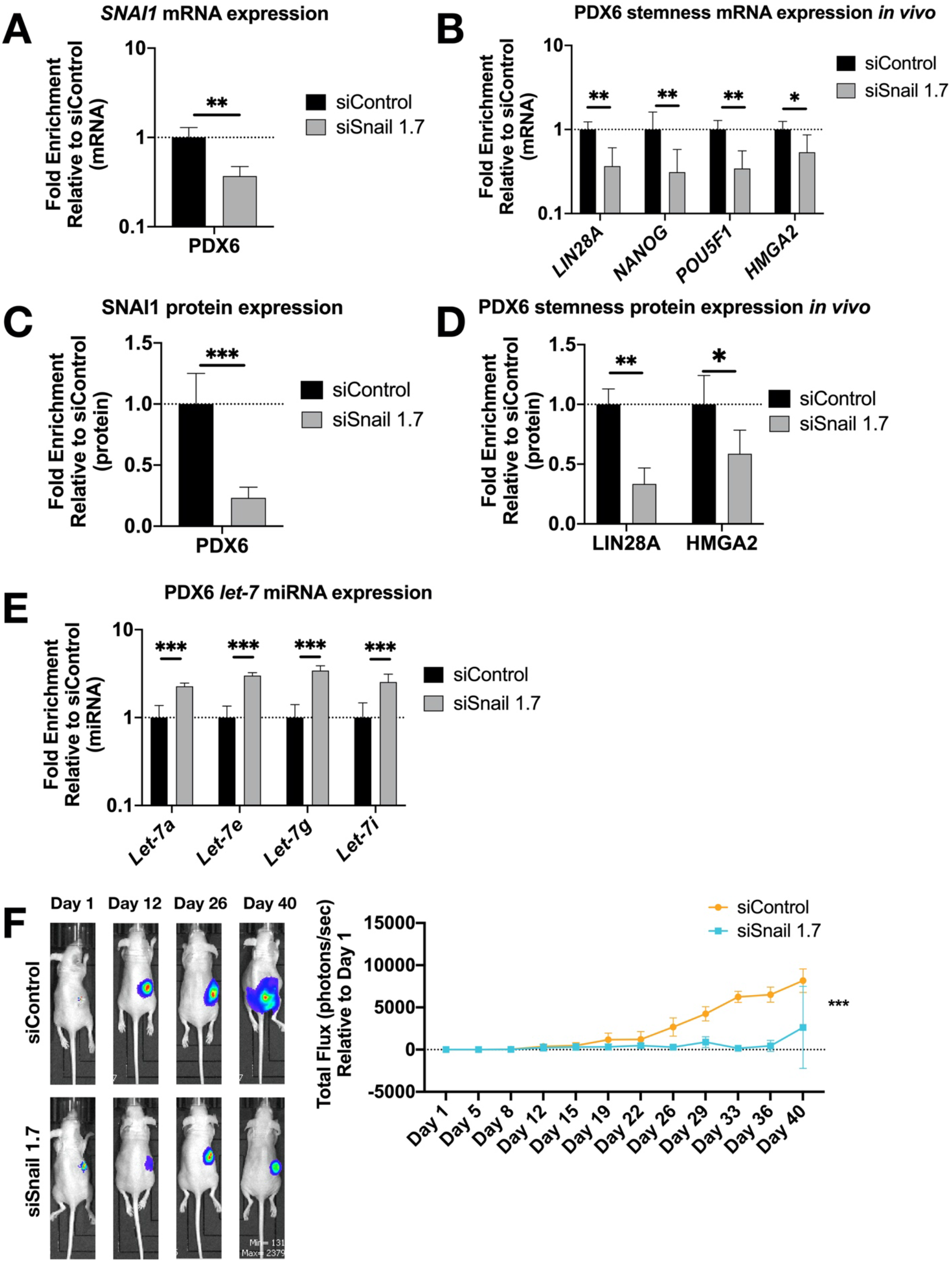
*SNAI1* knockdown *in vivo* reduces stemness gene expression and tumor burden. HA-MSN were used to deliver siRNA (siSnail and siControl) via IV injection to orthotopic PDX *in vivo*. Tumor samples were harvested and analyzed at necropsy. Levels of control group (cells treated with siControl) were normalized to 1; note that values for RT-qPCR are shown on a log scale. A,B. RT-qPCR analysis for mRNA expression of *SNAI1* (A) and stemness markers (B, *LIN28A, NANOG, POU5F1*, and *HMGA2*) in tumors. C,D. The quantification of Western blot analysis for protein expression of SNAI1 (C) and stemness markers (D, LIN28A and HMGA2) in tumors. E. RT-qPCR analysis for *let-7* miRNA (*let-7a, let-7e, let-7g* and *let-7i*) expression in tumors. F. Left panel: Representative images of xenograft mice. siControl (upper) and siSnail knockdown (lower). Right panel: Quantitation of bioluminescence at primary sites over six weeks. X axis, days; Y axis, total flux in photons/second relative to day 1.

### *SNAI1* binds *let-7* promoters resulting in *let-7* repression

We sought to establish whether *SNAI1* acts to directly repress let-7 transcription. *SNAI1* binds promoters of *let-7* in fibroblasts, and binding increases upon *SNAI1* overexpression^17^. To examine whether this same association can be observed in cancer cells, we carried out ChIP assays to determine binding of *SNAI1* to the promoter region of various *let-7* family members, as defined by previous studies^46–51^. The *let-7i* promoter is diagrammed in Figure 6A;^50,51^ the promoter region locations and the E-box (CANNTG) locations studied are listed in Supplementary Table 1. At baseline, we observed that *SNAI1* bound *CDH1* (used as a positive control) and *let-7* promoters to a greater extent in OVCAR8, the cell line with higher *SNAI1* expression, than in OVSAHO^56^ (Supplementary Figure 11A). We also assessed binding upon EMT induction by TGFB1 in MCF-7 cells and detected an increased level of *let-7i* and *miR-98* promoter binding compared to the control group (Supplementary Figure 11B). These data demonstrate *SNAI1* binding to *let-7* promoter regions in cancer cells tested.

**Figure 6.**
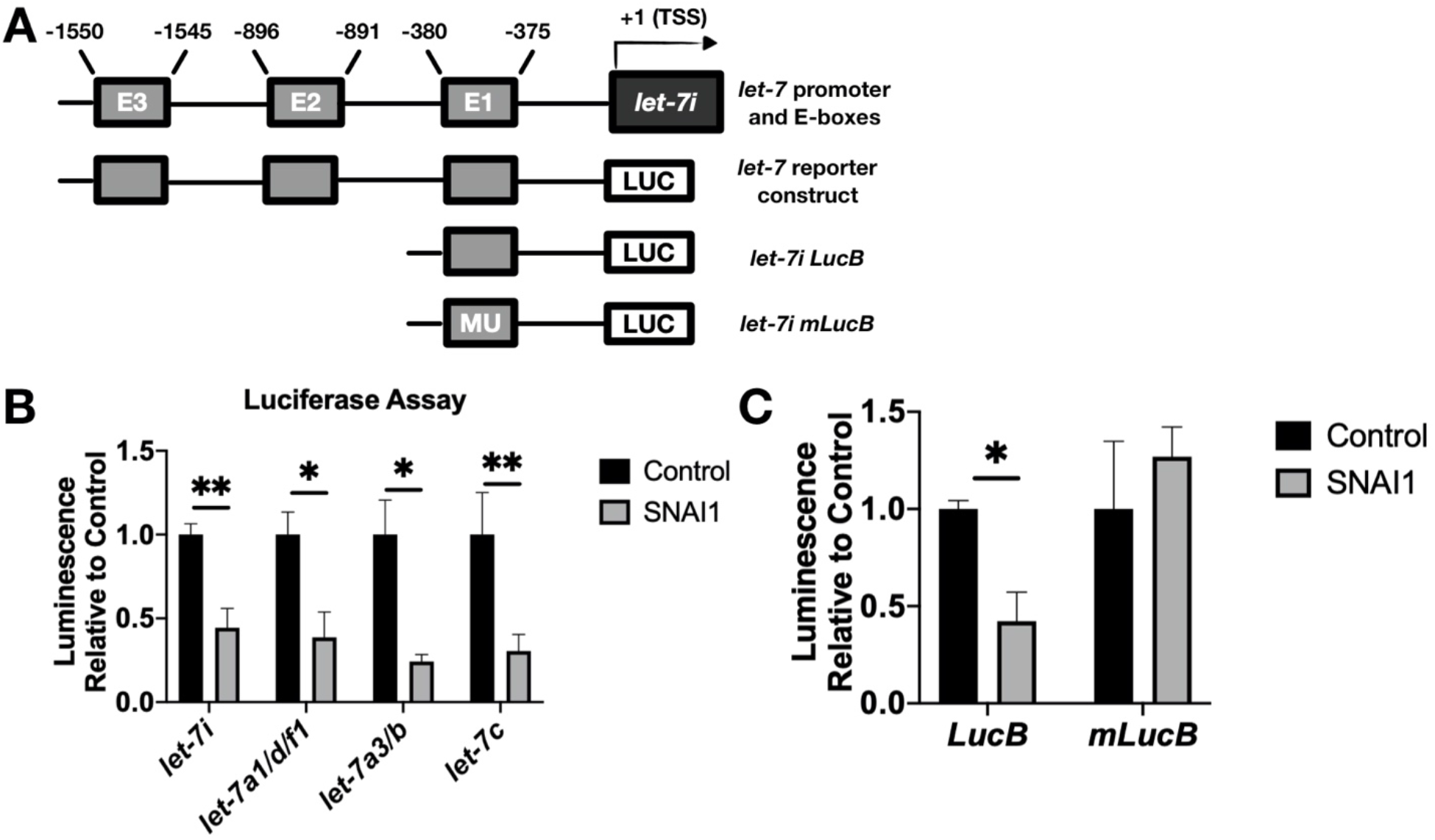
*SNAI1* represses *let-7* promoters. A. Schematic representation of the promoter region of *let-7i* (upper) and reporter constructs used in luciferase assays (lower diagrams). E1, E2, E3: E-boxes (sequence: CANNTG); MU: mutated E-boxes; TSS: transcription start site B. For luciferase assays, HEK293T cells were co-transfected with two plasmids: 1) *let-7* promoter luciferase (*let-7i, let-7a1/d/f1, let-7a3/b, let-7c*), and 2) either *SNAI1* (constitutively expressed, gray bars) or empty vector (black bars). Luminescence activity was measured 48 hours thereafter. C. HEK293T cells were co-transfected with either *let-7i* lucB or *let-7i* mlucB with or without *SNAI1*. Luminescence was measured 24 hours later.

To test the functional result of *SNAI1* binding to *let-7* promoters, luciferase assays were used as a reporter for *let-7* promoter activity via bioluminescence. We used let-7 promoter luciferase constructs as shown in Figure 6A (bottom diagram; see Supplementary Table 1). This enabled us to detect the effect of *SNAI1* on *let-7i, let7a1/d/f1, let-7a-3*, and *let-7c* promoter activity. Co-transfection with *let-7* promoter luciferase and *SNAI1* (constitutively expressed), as compared with empty vector, resulted in a reduction in bioluminescence (Figure 6B), confirming the repression of *let-7* promoter activity. A *let-7i* promoter luciferase mutated to remove E-box one was not inhibited by Snail (Figure 6C). These results demonstrate that *SNAI1* binding to *let-7* promoters directly represses *let-7* transcription.

## Discussion

*Let-7’s* major roles in maintenance of differentiation make it a key player in both development and cancer^13,14^. Loss of *let-7* is a major component of the loss of differentiation seen in many cancers, and significantly correlates with poor prognosis^13,16,18,19^. Studies of stem cell reprogramming linked *let-7* repression with a transcription factor that induces EMT, *SNAI1*^17^. In the present study, we examined the role of *let-7* in cancer cells and its connection to *SNAI1*. When cells from breast (MCF-7), pancreatic (PANC-1) and ovarian (OVCAR8, OVSAHO) cancer were treated with EMT-inducing agents (TGFB1 or EGF), increases in EMT factors including *SNAI1*, increases in stemness markers, and decreases in *let-7* could be detected. This positive association between *SNAI1* and stemness, as well as the negative association between *SNAI1* and *let-7*, were confirmed when *SNAI1* itself was overexpressed through viral transduction or knocked down by siRNA.

One of the goals of this investigation was to understand the molecular mechanisms by which *SNAI1* exerts its pro-stemness effects. The effect of *SNAI1* on *let-7* levels, and its direct binding to several *let-7* family member promoter regions, were detected using ChIP and luciferase assays, providing evidence that *SNAI1* binds *let-7* promoters and directly represses its expression, leading to an increase in stemness in cancer cells. Although EMT has been linked to stemness, few insights into downstream mechanisms have been generated. One downstream effector of *SNAI1* and other EMT programs is the transcription factor FOXC2 via the serine/threonine kinase p38, thus linking EMT and stem cell traits^35^. Another avenue by which *SNAI1* exerts stemness is via repression of miR-34 via effects on WNT signaling, NOTCH, and CD44^58^. Our results provide evidence for the *SNAI1/let-7* axis as another crucial mechanism by which EMT exerts pro-stemness roles. These results point to *SNAI1* as a stem cell-directed target for therapy.

*SNAI1* may be a particularly apt target in the goal of eliminating CSC because of its role in the stabilization of the hybrid epithelial-mesenchymal state^34,59^. OVCAR8 parental cells showed the highest level of stemness markers (*LIN28A, NANOG, POU5F1* and *HMGA2*, along with a high level of epithelial marker *CDH1* and mesenchymal markers *SNAI1* and *VIM* (Supplementary Figure 1A), suggesting its hybrid EMT status. *SNAI1* is highly expressed in all of the cell types examined here (Supplementary Figure 1A), and further studies will determine whether *SNAI1*-dependent *let-7* repression plays a role in the hybrid state.

*SNAI1* inhibition via transfection, viral delivery, or genetic deletion has been shown to reduce invasion, proliferation, chemoresistance, and other components of the stemness phenotype^56,60,61^. However, because these approaches cannot be considered for use in patients, other approaches such as nanoparticle-mediated delivery must be developed. Small RNAs can be efficiently loaded onto MSNs, which protect the oligonucleotides from degradation, are enriched in tumors due to leaky vasculature, and are taken up into cells by pinocytosis and as such function as a transfection reagent^42^. Their large surface area and pore structure make them ideal for drug delivery^40^. MSNs are a promising delivery agent for RNAi *in vivo*^41,45,62^. Considering this potential, and with the goal of clinical relevance, we used MSN to knock down *SNAI1. SNAI1* downregulation could be detected on both RNA and protein levels, emphasizing the utility of MSN for siRNA delivery. We extended these results to *in vivo* experiments where we knocked down *SNAI1* in our orthotopic PDX model^56^. We achieved >75% knockdown of SNAI1 protein in tumors *in vivo*, and importantly tumor *let-7* levels increased 2-3 fold, consistent with *SNAI1*-mediated repression of *let-7 in vivo*. In parallel, expression of stem cell markers *LIN28A, NANOG, POU5F1*, and *HMGA2* decreased, consistent with a shift away from the stem cell phenotype, demonstrating that targeting *SNAI1* is sufficient to reduce stemness. Further studies will determine if these changes lead to reduced metastasis or delayed recurrence.

Although these studies provide important insights into the mechanism for loss of *let-7* and thus the destabilization of the differentiated state, we do not address the question of the origin of CSC. Rather, we suggest that any cell of origin, in order to take on cancer stem cell characteristics, will lose *let-7*. Like differentiated cells, adult stem cells express high levels of *let-7*^63,64^, therefore *let-7* loss via transcriptional, post-transcriptional, or epigenetic regulation is required even in the case that adult stem cells are the cell of origin. In the absence of *LIN28A*, transcriptional repression of *let-7* could tip the balance in favor of stemness. The mechanism by which *let-7* is lost is thus germane to cancer stem cell biology regardless of whether normal stem cells or differentiated cells are the cells of origin. Our finding that *SNAI1* transcriptionally represses *let-7* adds even more weight to *SNAI1* as a therapeutic target: blocking *SNAI1*, in addition to inhibiting invasion and migratory ability, is expected to restore *let-7* by increasing its transcription. We predict that *SNAI1*-mediated *let-7* repression could be an important mechanism of cancer stemness in a wide variety of carcinoma cells.

## Acknowledgments

We thank Gottfried Konecny, Carlotta Glackin, Nathan Wall, and Eileen Brantley for cell lines, Jacqueline Coats for input on statistical analyses, members of the Perry lab for assistance with dynamic light scattering measurements of MSN, and members of the Unternaehrer lab for helpful discussions.

## Further Disclosures

### Data Availability Statement

Data will be made available from the corresponding author upon reasonable request

## List of abbreviations used

CSC: Cancer stem-like cells
ChIP: chromatin immunoprecipitation
DsiRNA: Dicer substrate small inhibitory RNA
EGF: Epidermal growth factor
EMT: Epithelial-mesenchymal transition
HA: hyaluronic acid
HGSOC: High grade serous ovarian carcinoma
IP: Immunoprecipitation
LLU: Loma Linda University
MSN: Mesoporous silica nanoparticles
PEI: polyethylenimine
PDX: Patient-derived xenograft
RT-qPCR: Reverse-transcription quantitative PCR
TGFB1: Transforming growth factor beta-1

