## Supplementary Figures and Tables for "The epithelial-mesenchymal transcription factor *SNAI1* represses transcription of the tumor suppressor miRNA *let-7* in cancer"

**Running title:**
*SNAI1* represses *let-7*

**Keywords:** epithelial-mesenchymal transition, stem cells, ovarian cancer, transcriptional regulation, miRNA, orthotopic patient-derived xenografts

**Supplementary Legends**

**Supplementary Figure 1.**

A. EMT status and stemness gene expression level comparison in parental cell lines. mRNA expression levels of epithelial (*CDH1*), mesenchymal (*SNAI1, VIM* and *CDH2*), and stemness markers (*LIN28A, NANOG, POU5F1*, and *HMGA2*, left to right) were analyzed using RT-qPCR. The expression levels were normalized to levels in NCCIT embryonal carcinoma cells (positive control for pluripotency gene expression), indicated as the dotted line.

B. TGFB1 does not induce EMT in OVSAHO or OVCAR8. mRNA expression levels of epithelial (*CDH1*) and mesenchymal (*SNAI1, VIM and CDH2*) factors (from left to right) were analyzed using RT-qPCR after OVCAR8 (left) and OVSAHO (right) were treated with TGFB1. The expression levels of markers in control groups (cells treated with vehicle control) were normalized to 1, indicated as the dotted line.

**Supplementary Figure 2.**

A. Western blot analysis for protein expression of SNAI1 in MCF-7, PANC-1, OVCAR8, and OVSAHO (samples from Fig. 1C). α/β-TUBULIN was used as a loading control.

B. Western blot analysis for protein expression of HMGA2 in OVSAHO (samples from Fig. 1D). α/β-TUBULIN was used as a loading control.

**Supplementary Figure 3**. ***SNAI1 siRNA screening.*** Twelve DsiRNAs were screened for their effectiveness at SNAI1 knockdown in HEK293T cells. Lipofectamine RNAiMax (Life Technologies) was used for transfection of 1, 3, or 10nM DsiRNAs. After 24h, levels of *SNAI1* RNA using a 5' (*SNAI1*-1) and a 3' (*SNAI1*-2) primer pair were determined with RT-qPCR. As a positive control, HPRT was targeted; siControl was used as a negative control. siSnail 1.7 was chosen as most effective in dose response for SNAI1 knockdown. siSnail 1.6 were also used for *in vitro* knockdown with similar results.

**Supplementary Figure 4.**

A. Western blot analysis for protein expression of SNAI1 in MCF-7, PANC-1, OVCAR8, and OVSAHO (samples from Fig. 2C). α/β-TUBULIN was used as a loading control.

B. Western blot analysis for protein expression of HMGA2 in OVCAR8 and OVSAHO (samples from Fig. 2D). α/β-TUBULIN was used as a loading control.

C. Phase contrast images (3 representative images) of spheroids formed from MCF-7, PANC-1, OVCAR8 and OVSAHO are presented; in each panel, the spheroids formed from control group (cells transduced with pWZL-Empty) are presented on the left, those from pWZL-Snail group are on the right (quantified in Fig. 2E).

**Supplementary Figure 5.** ***Higher level of SNAI1 is associated with larger size of spheroids and higher number of secondary spheroids formed.***

A. The quantification of spheroid size (unit: $microns^2$) for OVCAR8 in *SNAI1* overexpression (left), SNAI1 knockdown (middle) and PDX6 cells in *SNAI1* knockdown *in vitro* (right);

B. Phase contrast images of spheroids (after passage) for EMT induction via TGFB1/EGF (top panel), *SNAI1* retroviral overexpression (middle panel) and *SNAI1* knockdown (lower panel). In each panel, data of MCF-7, PANC-1, OVCAR8 are shown (left to right); the quantification of number of spheroids per 3000 cells is presented in the graph to the right of the pictures.

**Supplementary Figure 6.** ***Let-7i overexpression rescues the increase in stemness resulting from SNAI1 overexpression.***

A. For the rescue experiment, cells transduced with pWZL-Snail were also transfected with control mimic or *let-7i* mimic afterwards; miRNA expression levels of *let-7i* were analyzed using RT-qPCR. The expression levels of *let-7i* in control groups (cells transduced with pWZL-Snail then transfected with Mimic Control) were normalized to 1.

B. RT-qPCR analysis for stemness markers (*LIN28A, NANOG, POU5F1*, and *HMGA2*) in MCF-7, PANC-1, OVCAR8 and OVSAHO are presented, note only the cells transduced with pWZL-Empty then transfected with Mimic Control were normalized to 1.

C. Phase contrast images of spheroids for MCF-7, PANC-1, OVCAR8 and OVSAHO (left to right); in each panel, pictures from control mimic group are shown on the left, those from *let-7i* mimic group on the right.

D. The quantification of number of spheroids per 3000 cells formed from MCF-7, PANC-1, OVCAR8 and OVSAHO in the rescue experiment.

**Supplementary Figure 7.**

A. Western blot analysis for protein expression of SNAI1 in MCF-7, PANC-1, OVCAR8, and OVSAHO (quantified in Fig. 3C). α/β-TUBULIN was used as a loading control.

B. Western blot analysis for protein expression of HMGA2 in PANC-1 and OVSAHO (quantified in Fig. 3D). α/β-TUBULIN was used as a loading control.

C. Phase contrast images (3 representative images) of spheroids formed from MCF-7, PANC-1, OVCAR8 and OVSAHO are presented; in each panel, the spheroids formed from control group (cells transduced with siControl) are presented on the left, those from siSnail 1.7 group are on the right (quantified in Fig. 3E).

**Supplementary Figure 8. Knockdown of *SNAI1* with a second DsiRNA.**

A. mRNA expression level of *SNAI1* in MCF-7, PANC-1, OVCAR8 and OVSAHO after treatment of DsiRNA 1.6 was analyzed using RT-qPCR. The expression level of *SNAI1* in control groups (cells treated with siControl) was normalized to 1, indicated as the dotted line. B. Phase contrast images of spheroids formed from siControl group (left, same images as Figure 3E) and siSnail 1.6 group (right). The quantification of number of spheroids formed per 3000 cells is shown below.

**Supplementary Figure 9.**

A. Western blot analysis for protein expression of SNAI1 and HMGA2 in PDX6 *in vitro* (quantified in Fig. 4C and D). α/β-TUBULIN was used as a loading control. Triplicates of each group are presented.

B. Phase contrast images (3 representative images) of spheroids formed from PDX6 are presented (quantified in Fig. 4E).

**Supplementary Figure 10.**

A. Western blot analysis for protein expression of SNAI1 HMGA2 and LIN28A in PDX6 *in vivo* (quantified in Fig. 5C and D). α/β-TUBULIN was used as a loading control.

B. The average tumor weights in mice treated with siControl (black) vs. siSnail (gray) *in vivo*.

**Supplementary Figure 11. SNAI1 binds to *let-7* promoters**

Chromatin immunoprecipitation (ChIP) analysis of SNAI1 binding to *let-7* promoter regions. SNAI1 binding *to let-7a3-b, let-7c, let-7g, let-7i, mir98* was measured as fold enrichment relative to IgG negative control. *CDH1* was used as an internal positive control.

A. Parental cell lines of OVCAR8 and OVSAHO were collected and their levels of SNAI1 binding to *let-7* promoter were measured by ChIP.

B: MCF-7 cells were treated with TGFB1. Shown are cells treated with vehicle control vs TGFB1. The table above shows the exact fold enrichment for each primer.

**Supplementary Table 1.** *Let-7* promoter region length, locations and E-box (CANNTG) locations

**Supplementary Table 2.** Sequences of primers used in RT-qPCR

**Supplementary Table 3.** Sequences of primers used in ChIP

**Supplementary Table 4.** Sequences of siRNA oligonucleotides

**Supplementary Figure 1**

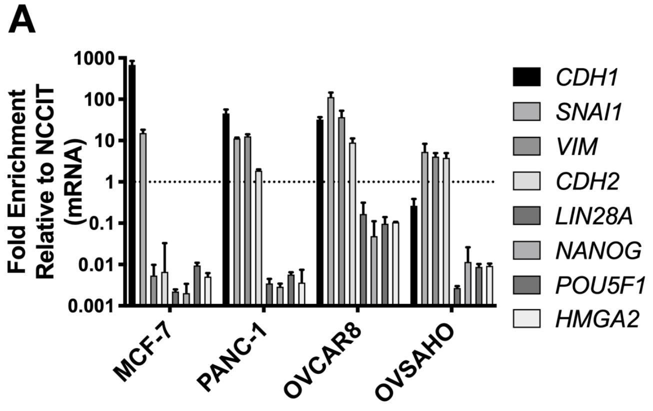


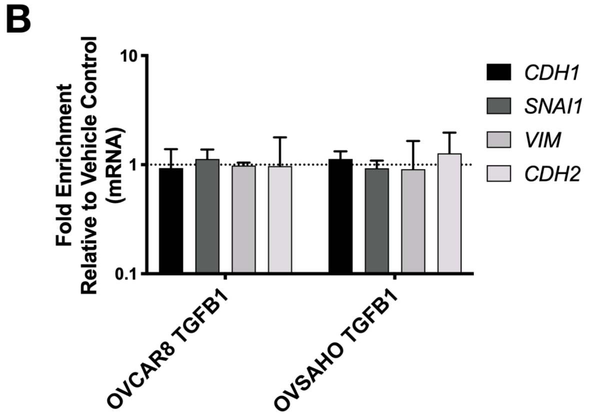

**Supplementary Figure 2**

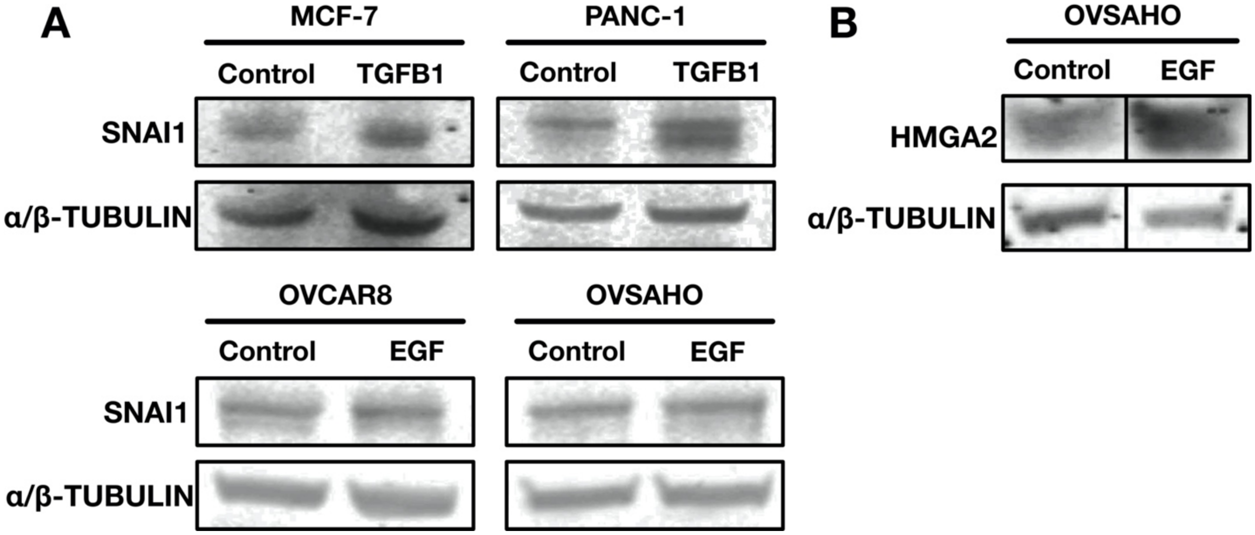

#### Supplementary Figure 3

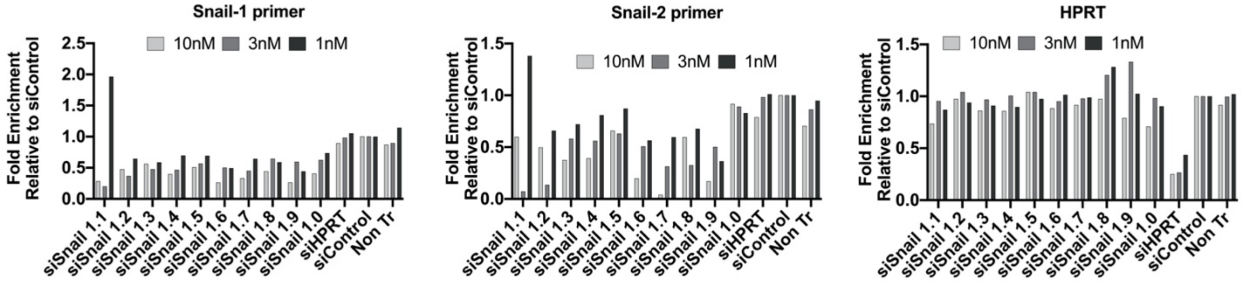

#### Supplementary Figure 4

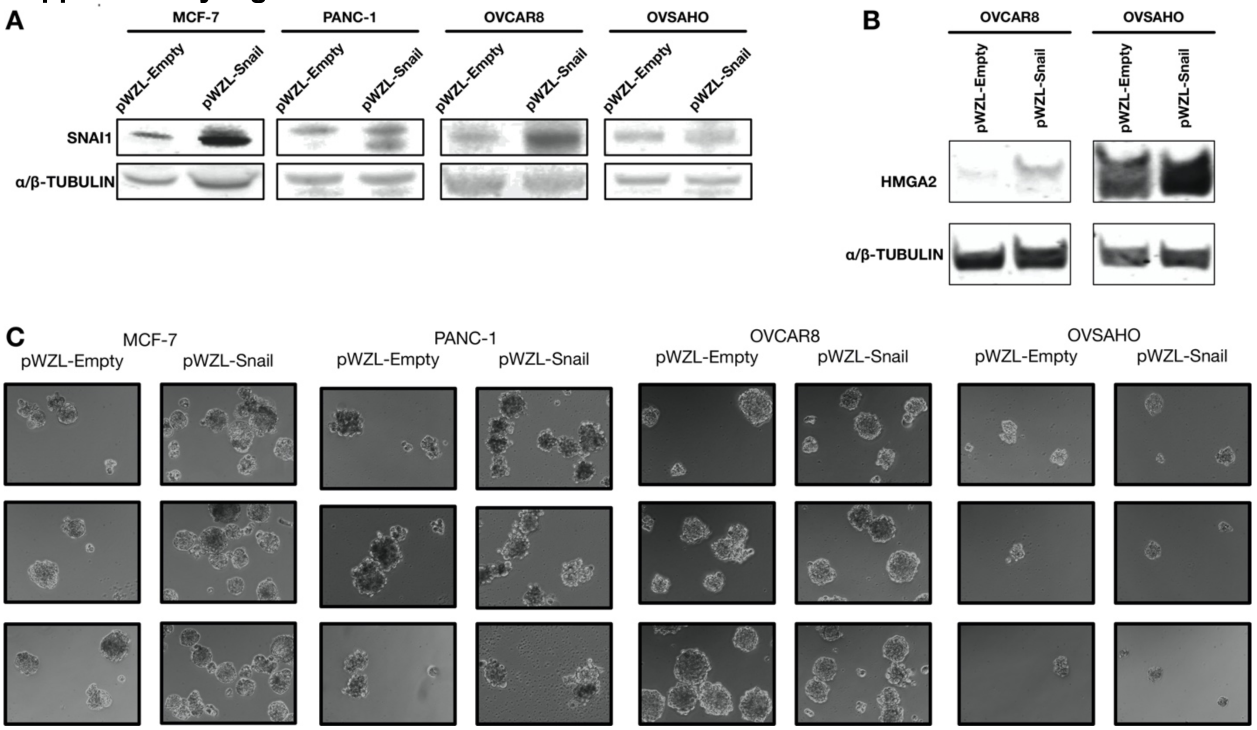

### Supplementary Figure 5

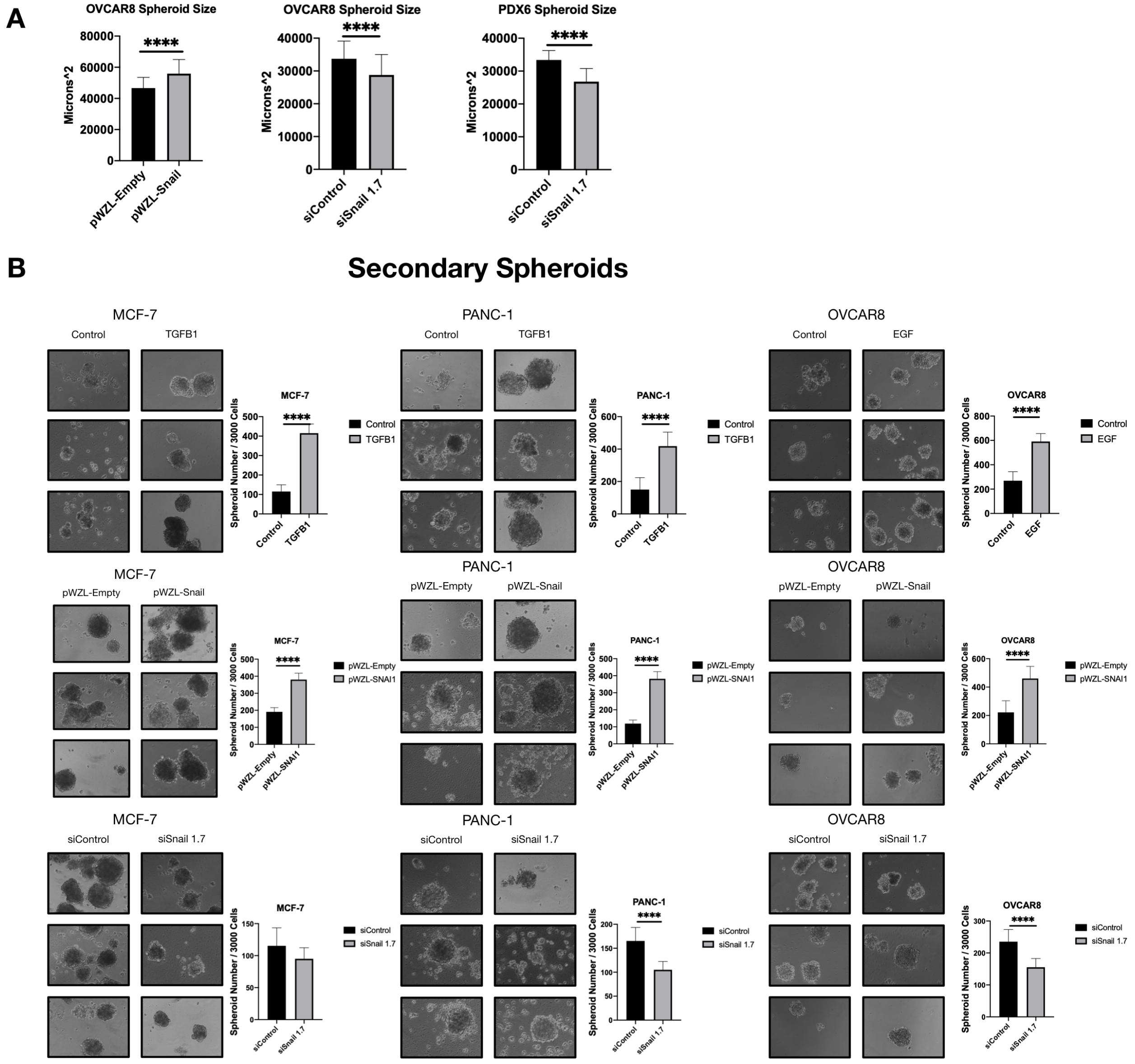

#### Supplementary Figure 6

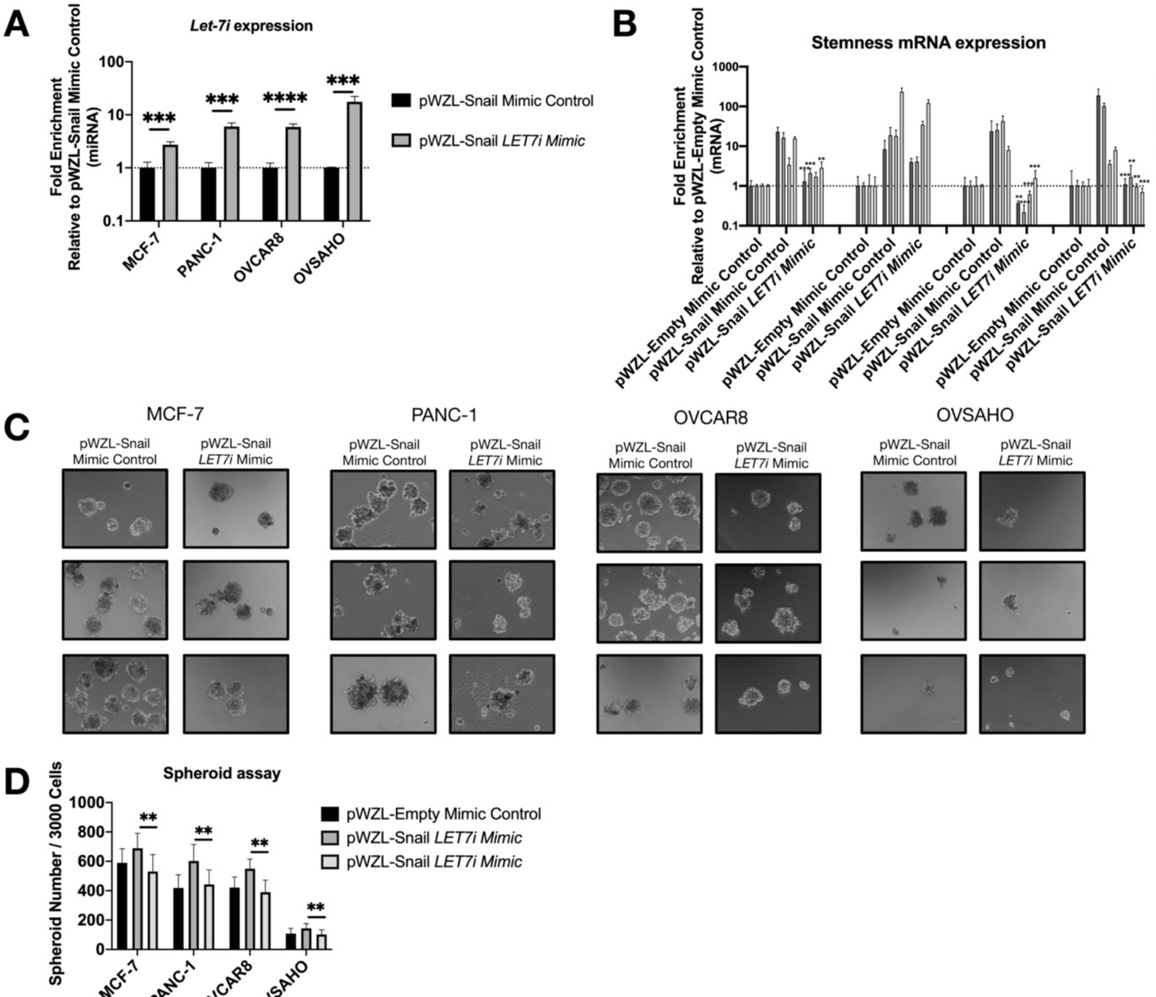

#### Supplementary Figure 7

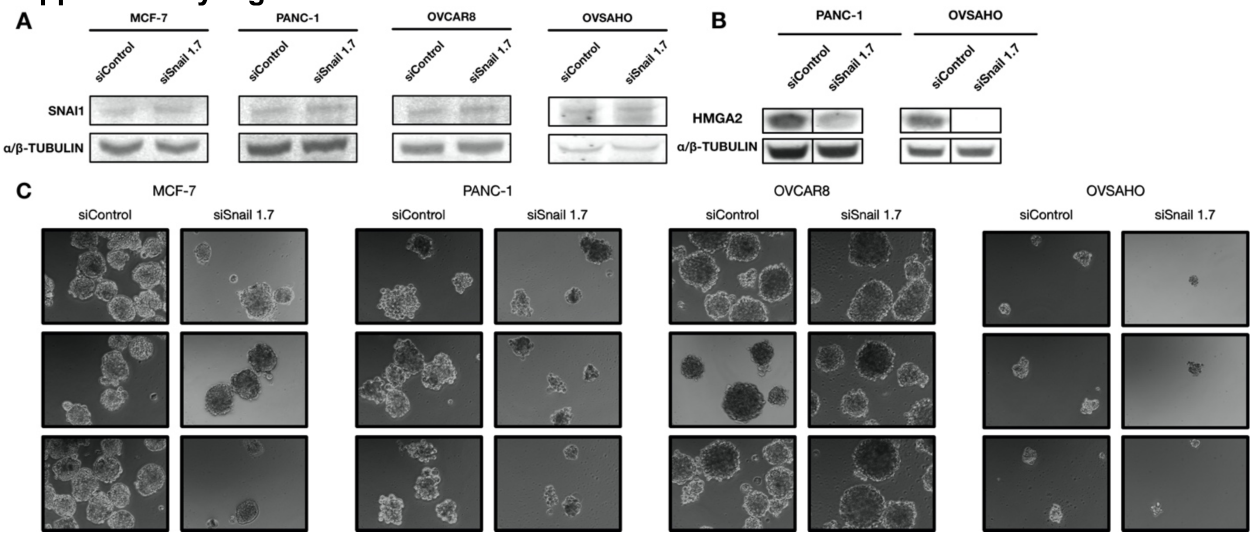

#### Supplementary Figure 8

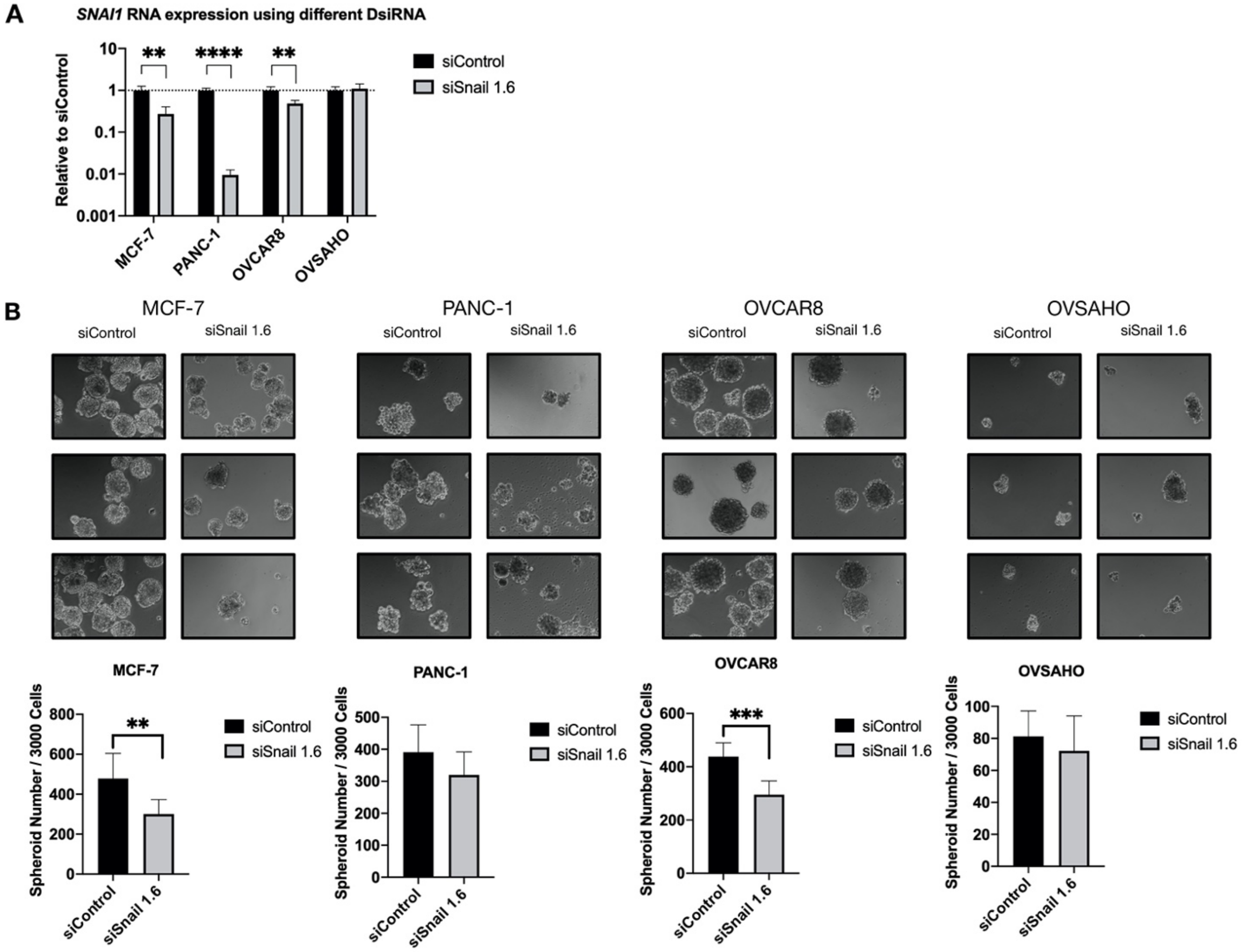

**Supplementary Figure 9**

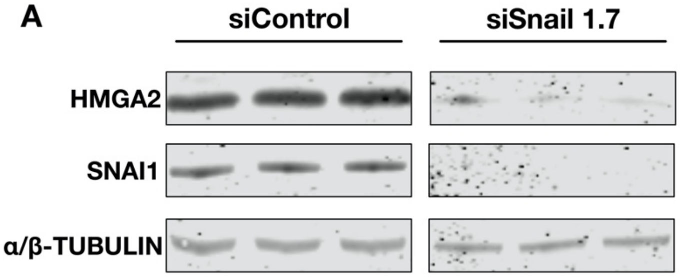


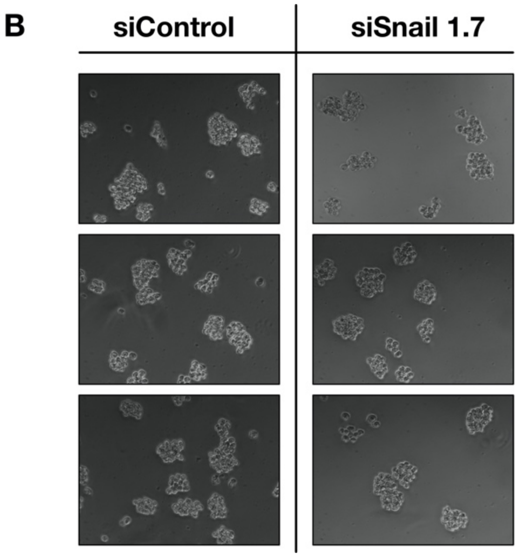

Supplementary Figure 10

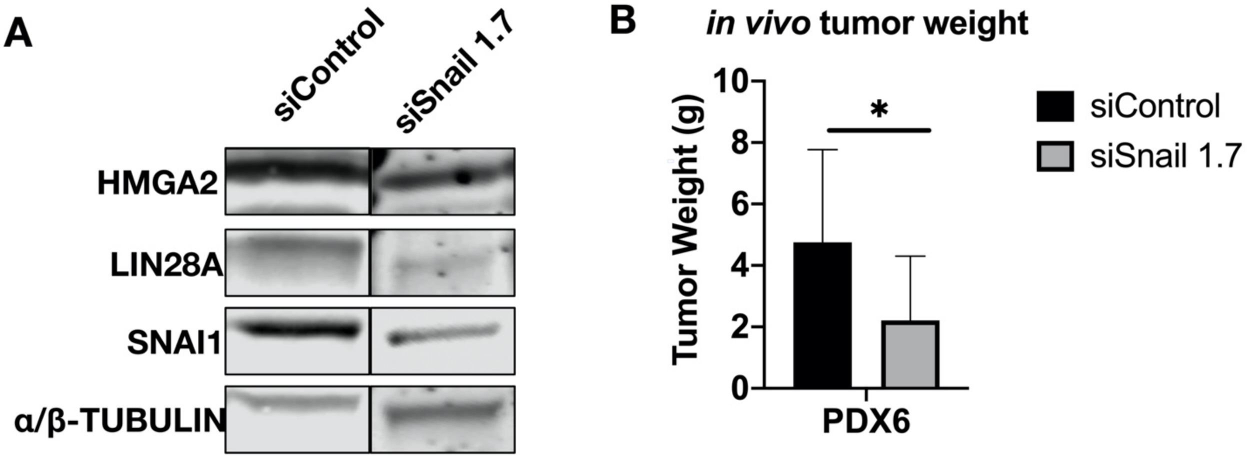

#### Supplementary Figure 11

A

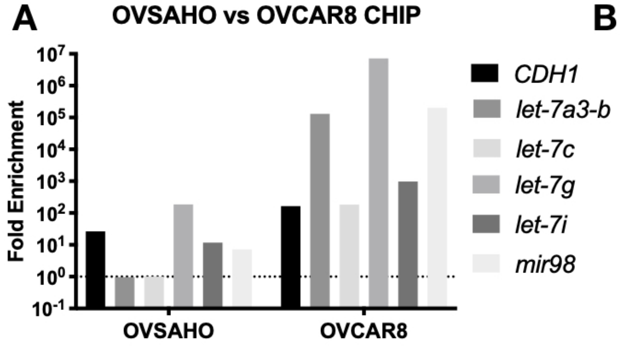


B

| | *CDH1* | *let-7a3-b* | *let-7c* | *let-7g* | *let-7i* | *mir98* |
|---|---|---|---|---|---|---|
| Control | 6.05732488 | 0.03138349 | 0.00449943 | 0.00025097 | 0.60049700 | 0.0000011922 |
| TGFB1 | 358.25656400 | 3.09103324 | 1.00000000 | 1.00000000 | 403.67517800 | 523.7606820000 |

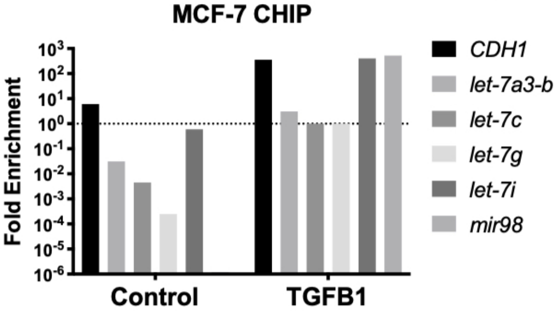

**Supplementary Table 1**

| *let-7* member | E-box starting site: bp upstream of TSS | Promoter region starting site to ending site: bp upstream of TSS |
|---|---|---|
| *let-7a1-f1*[1] | -8924, -8431 | -9775 to -8286 |
| *let-7a3-b*[2] | -1688, -1581, -1302, -1234 | -1960 to -917 |
| *let-7c*[3] | -1783, -1526, -1501, -1326, -1279, -640, -481, -247, -197 | -1804 to 0 |
| *let-7g*[4] | -11259, -9824, -9194, -9045 | -11397 to -9000 |
| *let-7i*[5] | -1545, -891, -375 | -2303 to -39 |
| *mir98*[6] | -4479, -2749, -2217, -1249 | -5032 to - 536 |

**Supplementary Table 2**

| primers | Forward | Reverse |
|---|---|---|
| *ACTB* | 5' - TGAAGTGTGACGTGGACATC - 3' | 5' - GGAGGAGCAATGATCTTGAT - 3' |
| *SNAI1 Snail-1* | 5' - CACTATGCCGCGCTCTTTC - 3' | 5' - GGTCGTAGGGCTGCTGGAA - 3' |
| *SNAI1 Snail-2* | 5’ - GGC TGC TAC AAG GCC AT - 3’ | 5’ - GCA CTG GTA CTT CTT GAC ATC T - 3’ |
| *NANOG* | 5' - CAAAGGCAAACAACCCACTT - 3' | 5' - TCTGCTGGAGGCTGAGGTAT - 3' |
| *LIN28A* | 5' - GAGCATGCAGAAGCGCAGATCAAA - 3' | 5' - TATGGCTGATGCTCTGGCAGAAGT - 3' |
| *POU5F1* | 5' - AAGCGATCAAGCAGCGACTAT - 3' | 5' - GGAAAGGGACCGAGGAGTACA - 3' |
| *HMGA2* | 5' - GCAGCCGTCCACTTCAG - 3' | 5' - TCTTCCCCTGGGTCTCTTAG - 3' |
| *HPRT* | 5’ - GGCTTATATCCAACACTTCG GGG - 3’ | 5’ - GACTTTGCTTTCCTTGGTCAG - 3’ |

**Supplementary Table 3**

| ChIP primers | Forward | Reverse |
|---|---|---|
| *Let-7a3-b* | 5' - GTCATCCTCATGCTACACTCTG - 3' | 5' - AAAGGAGGAGGGAGGGAAA - 3' |
| *Let-7c* | 5' - ACTCAGATGCTCCTCCTAGAC - 3' | 5' - CTCCTCCTGATTTCACTTCCTTT - 3' |
| *Let-7g* | 5' - GGTACTCGATCTGTCCTTGTTG - 3' | 5' - GGAAGAGTTGCACTATCCTGTT - 3' |
| *Let-7i* | 5' - GGAGGGATTTCAGGCAAGAT - 3' | 5' - ATGAAGGGACCCAATCCTTC - 3' |
| *Mir98* | 5' - TGACTGTGTCTGCCGTAATG - 3' | 5' - AGTGTGACTGTTGAGTGAGAAG - 3' |
| *CDH1* | 5’ – GTTTGTTTGTTTTCCAAC – 3’ | 5’ – GCAAGGCTATGTCTCAAAAG – 3’ |

**Supplementary Table 4**

| RNA oligonucleotides | Sense (5' - 3') | Antisense (5' - 3') |
|---|---|---|
| siSnail 1.1 | rArGrArUrGrUrCrArArGrArArGrUrArCrCrArGrUrGrCrCAG | rCrUrGrGrCrArCrUrGrGrUrArCrUrUrCrUrUrGrArCrArUrCrUrGrA |
| siSnail 1.2 | rUrGrUrGrArCrUrArArCrUrArUrGrCrArArUrArArUrCrCAC | rGrUrGrGrArUrUrArUrUrGrCrArUrArGrUrUrArGrUrCrArCrArCrC |
| siSnail 1.3 | rUrCrArArCrUrGrCrArArArUrArCrUrGrCrArArCrArArGGA | rUrCrCrUrUrGrUrUrGrCrArGrUrArUrUrUrGrCrArGrUrUrGrArArG |
| siSnail 1.4 | rGrCrGrArGrCrUrGrCrArGrGrArCrUrCrUrArArUrCrCrAGA | rUrCrUrGrGrArUrUrArGrArGrUrCrCrUrGrCrArGrCrUrCrGrCrUrG |
| siSnail 1.5 | rGrGrCrCrUrArGrCrGrArGrUrGrGrUrUrCrUrUrCrUrGrCGC | rGrCrGrCrArGrArArGrArArCrCrArCrUrCrGrCrUrArGrGrCrCrGrU |
| siSnail 1.6 | rCrUrArCrArGrGrArCrArArArGrGrCrUrGrArCrArGrArCTC | rGrArGrUrCrUrGrUrCrArGrCrCrUrUrUrGrUrCrCrUrGrUrArGrCrU |
| siSnail 1.7 | rGrCrCrArArUrGrCrUrCrArUrCrUrGrGrGrArCrUrCrUrGTC | rGrArCrArGrArGrUrCrCrCrArGrArUrGrArGrCrArUrUrGrGrCrArG |
| siSnail 1.8 | rArCrUrGrUrGrArGrUrArArUrGrGrCrUrGrUrCrArCrUrUGT | rArCrArArGrUrGrArCrArGrCrCrArUrUrArCrUrCrArCrArGrUrCrC |
| siSnail 1.9 | rCrArArGrArUrGrCrArCrArUrCrCrGrArArGrCrCrArCrACG | rCrGrUrGrUrGrGrCrUrUrCrGrGrArUrGrUrGrCrArUrCrUrUrGrArG |
| siSnail 1.0 | rUrUrCrCrCrGrGrGrCrArArUrUrUrArArCrArArUrGrUrCTG | rCrArGrArCrArUrUrGrUrUrArArArUrUrGrCrCrCrGrGrGrArArArC |
| siHPRT | rGrCrCrArGrArCrUrUrUrGrUrUrGrGrArUrUrUrGrArArATT | rArArUrUrUrCrArArArUrCrCrArArCrArArArGrUrCrUrGrGrCrUrU |
| siControl | rCrArUrArUrUrGrCrGrCrGrUrArUrArGrUrCrGrCrGrUrUAG | rCrUrArArCrGrCrGrArCrUrArUrArCrGrCrGrCrArArUrArUrGrGrU |
| siSnail 1.7 mod | mGmCrCmArAmUrGrCrUrCrAmUrCmUrGmGrGmArCrUrCrUrGmUC | rGrAmCrArGrArGrUmCrCmCrAmGrArUrGrArGrCrArUrUmGrGmCmAmG |
| siControl mod | mCmArUmArUmUrGrCrGrCrGmUrAmUrAmGrUmCrGrCrGrUrUmAG | rCrUmArArCrGrCrGmArCmUrAmUrArCrGrCrGrCrArArUmArUmGmGmU |

### Western Blot whole membrane used in manuscript.
### Supplementary Material for review

A

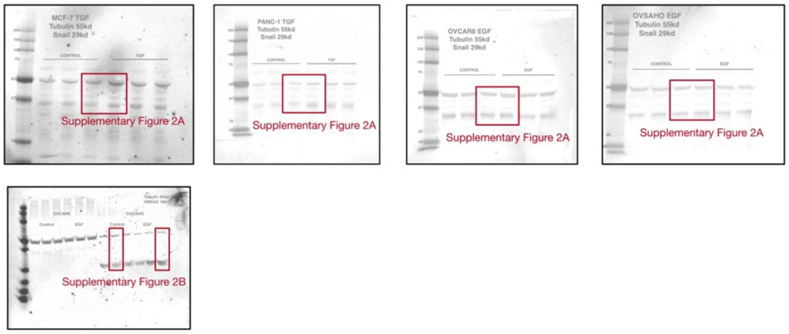


B

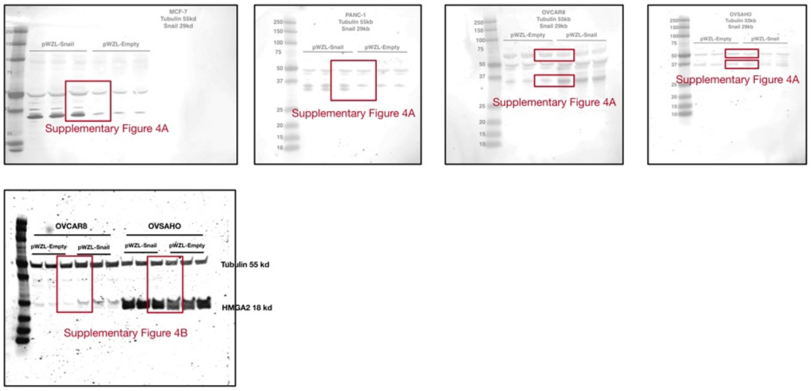


C

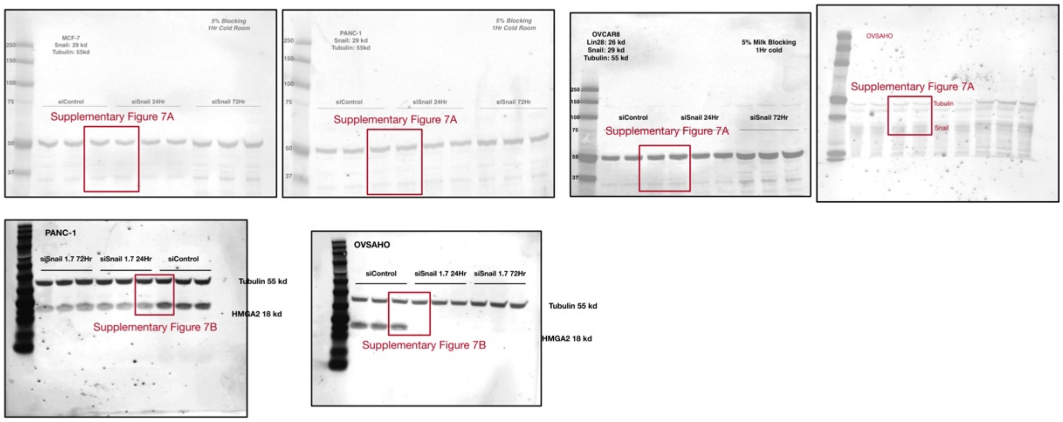


D

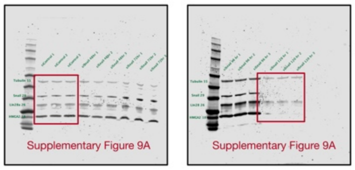


E

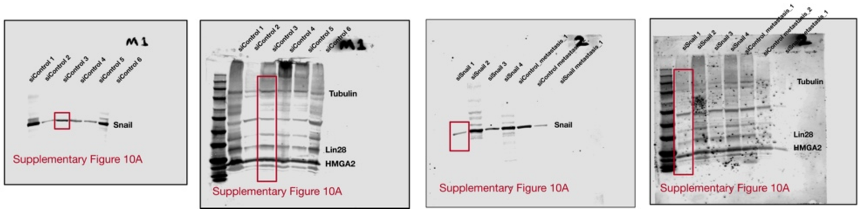

Supplementary Material for Review
These are the immunoblots that were cropped and used in the manuscript. The exact band used in manuscript are labeled.

A) Refers to Supplementary Figure 2. The Western blot results for SNAI1 in TGFB1/EGF treatments for MCF-7, PANC-1, OVCAR8 and OVSAHO are presented here from left to right (one cell line per blot). In each blot, the left three lanes were samples from control group, the right three lanes were samples from treatment group (upper panel). The blot for HMGA2 bands in OVCAR8 and OVSAHO after EGF treatment is presented in lower panel. The lanes from control and EGF group of OVCAR8 and OVSAHO are indicated in the blot. One lane from each group was picked and presented in the figure. The position of TUBULIN bands, HMGA2 bands and SNAI1 bands could be determined based on the molecular weight markers. Additionally, the signal for TUBULIN, HMGA2, LIN28 is visible in the 700 nm channel, while that for SNAI1 is visible in the 800 nm channel; this also applies to B, C, D and E.

B) Refers to Supplementary Figure 4. The Western blot results for SNAI1 in *SNAI1* overexpression samples for MCF-7, PANC-1, OVCAR8 and OVSAHO are presented from left to right (one cell line per blot). The result for HMGA2 in OVCAR8 and OVSAHO is presented in the lower panel. The lanes from pWZL-Empty and pWZL-Snail group are indicated in each blot. One lane from each group was picked and presented in the figure. The position of TUBULIN bands and SNAI1 bands could be determined based on the molecular weight markers.

C) Refers to Supplementary Figure 7. The Western blot results for SNAI1 in *SNAI1* knockdown samples for MCF-7, PANC-1, OVCAR8 and OVSAHO are presented from left to right (one cell line per blot). The Western blot results for HMGA2 in *SNAI1* knockdown samples for PANC-1 and OVSAHO are presented (one cell line per blot). The lanes from siControl; siSnail 1.7 24Hr and siSnail 1.7 48 Hr group are indicated in each blot. One lane from each group (siControl and siSnail 1.7 24Hr) was picked and presented in the figure. The position of TUBULIN bands and HMGA2 bands are indicated on the blot.

D) Refers to Figure 4. The Western blot results of PDX6 cells treated with siSnail 1.7 *in vitro*. From left to right were siControl samples, siSnail 1.7 48 hrs, siSnail 1.7 72 hrs (left blot), siSnail 1.7 96 hrs, siSnail 120 hrs (right blot), each in triplicate. The samples from siControl and siSnail 120 hrs were picked and presented in the figure. The position of TUBULIN bands, SNAI1 bands, LIN28 bands and HMGA2 bands are noted on the blots.

E) Refers to Figure 5. The Western blot results of PDX6 cells treated with siSnail 1.7 *in vivo*. The two blots on the left are identical blots scanned at 800 nm (left) and 700 nm (right) to visualize SNAI1 (left) and TUBULIN, LIN28A, HMGA2 (right). The same applies to the two blots on the right. In the two blots on the left, each lane represented a primary tumor sample from one mouse from siControl group. In the two blots on the right, the first four lanes each represented a primary tumor sample from one mouse from siSnail group. The last three lanes are metastasis tumor samples that are from siControl and siSnail (not presented in manuscript). The position of TUBULIN, SNAI1, LIN28, and HMGA2 bands are noted on the blots.
